# Curation of over 10,000 transcriptomic studies to enable data reuse

**DOI:** 10.1101/2020.07.13.201442

**Authors:** Nathaniel Lim, Stepan Tesar, Manuel Belmadani, Guillaume Poirier-Morency, Burak Ogan Mancarci, Jordan Sicherman, Matthew Jacobson, Justin Leong, Patrick Tan, Paul Pavlidis

**Author notes:** Corresponding author: Paul Pavlidis, 177 Michael Smith Laboratories, 2185 East Mall, University of British Columbia, Vancouver BC V6T1Z4.

## Abstract

Vast amounts of transcriptomic data reside in public repositories, but effective reuse remains challenging. Issues include unstructured dataset metadata, inconsistent data processing and quality control, and inconsistent probe-gene mappings across microarray technologies. Thus, extensive curation and data reprocessing is necessary prior to any reuse. The Gemma bioinformatics system was created to help address these issues. Gemma consists of a database of curated transcriptomic datasets, analytical software, a web interface, and web services. Here we present an update on Gemma’s holdings, data processing and analysis pipelines, our curation guidelines, and software features. As of June 2020, Gemma contains 10,811 manually curated datasets (primarily human, mouse, and rat), over 395,000 samples and hundreds of curated transcriptomic platforms (both microarray and RNA-sequencing). Dataset topics were represented with 10,215 distinct terms from 12 ontologies, for a total of 54,316 topic annotations (mean topics/dataset = 5.2). While Gemma has broad coverage of conditions and tissues, it captures a large majority of available brain-related datasets, accounting for 34% of its holdings. Users can access the curated data and differential expression analyses through the Gemma website, RESTful service, and an R package.

Database URL: https://gemma.msl.ubc.ca/home.html

## Introduction

As of June 2020, NCBI Gene Expression Omnibus (GEO) contains 97,379 transcriptomic studies, 71,233 (73%) of which are generated in human, mouse and rat (1). This vast resource presents a rich opportunity for data reuse and secondary analyses. For instance, many meta-analyses have been performed using published data, borrowing power across similar studies to draw conclusions with greater confidence (2–9); generation of co-expression networks (10), model generation for predicting gene expression (11), and the creation of derivative databases (12–14).

While the data in GEO is plentiful, in our experience, substantial effort in curating and reprocessing is necessary before high-quality secondary analyses can be performed. This effort is necessary because GEO was originally intended mainly to be an archive of published data (15). Accordingly, it was designed to accept as many types of data possible with low burden on data submitters, thus necessitating a relatively simple data model. While this may have been important to the rapid uptake of GEO as a key repository, it leaves researchers who want to reuse the data with some problems. These problems can be roughly grouped into affecting three aspects: metadata, microarray platform probe annotations, and the expression data itself.

In some datasets, the available metadata consist of a very brief description of the study and cryptically-named samples; in others, there is insufficient sample information to easily reconstruct the underlying study design, despite the MIAME standard GEO follows (16). This problem is exacerbated when, as is sometimes the case, the relevant publication is not linked to the dataset record. Regarding microarray platforms, different manufacturers have varying methods of associating probes to genes, and this inconsistency may result in lower comparability between datasets. Additionally, probe sequences for many microarray platforms were not submitted to GEO, thus the mapping of probes to genes cannot be substantiated, much less reproduced or updated, without referring to other sources of information. Lastly, expression data in GEO does not undergo any quality control aside from whatever the submitters might have performed, and it cannot be assumed that this was sufficient (17). Furthermore, while GEO stores raw expression data (e.g. Affymetrix CEL files, RNA-Sequencing FASTQ files etc.), the processed data is provided by the submitters, leaving consistency of processing approach a concern. For all of these reasons, instead of simply working directly with processed data and metadata downloaded from GEO, users are compelled to (or at least should) independently annotate, reprocess and perform quality control on the downloaded raw data before performing downstream analyses.

To help address these issues, we created Gemma (18), a bioinformatics system consisting of a database of curated gene expression studies sourced primarily from GEO, analysis pipelines, and accompanying website and web services. In this paper, we describe the data processing and curation pipelines in Gemma and the key design rationales involved, provide summaries and updates on the data contained in Gemma’s database, compare and contrast Gemma with other similar projects in the bioinformatics literature, and present a summary of notable Gemma features.

## Methods

### Data model

The Gemma data model was influenced by (but does not fully adopt) an early microarray data model, MAGE-OM (19), and some of our internal terminology carries its vestiges. Central entities in the Gemma data model are ExpressionExperiments (representing datasets of multiple samples, often corresponding to a single GEO Series (GSE)) and ArrayDesigns (transcriptome platforms, generally corresponding to a single GEO Platform (GPL); the same concept is adapted for RNA sequencing (RNA-seq)). ExpressionExperiments are made up of BioAssays (approximately corresponding to a GEO sample (GSM IDs)), which bring together a BioMaterial (the RNA sample) and an ArrayDesign. Note that GEO does not have a separate concept of a BioMaterial, but this concept is necessary to accommodate datasets where the same RNA sample was run on more than one platform (discussed further below under “Processing of expression data”). For each ExpressionExperiment, the expression data itself is stored at the level of CompositeSequences (platform elements: probes or probe sets for microarrays, genes for RNA-seq). CompositeSequences are mapped to GeneProducts (and thus Genes) via separate BioSequenceAssociation entities. The Gemma model has other entities to model sample annotations, publications, external database references (e.g. to GEO), data analyses, groups of datasets or genes, as well as security-related concepts such as users, full description of which is omitted here.

### The Gemma framework

Processing of large amounts of gene expression data into Gemma involves numerous steps. Major grouping of these steps include dataset selection, platform processing, expression data processing, metadata curation, and downstream analyses. Many of these steps are automated, with some human intervention and manual curation at key stages, described in the subsequent sections. Before delving further, we first lay out the decisions that influenced the framework’s design.

The main goal of Gemma’s data processing and curation pipelines is to provide a rich yet usable representation of a dataset; the representation can be described as having three parts. First, the expression data itself is needed, such that the data can be usefully represented as a single matrix where rows represent “platform elements” (genes, probes or probe sets) and columns represent RNA samples, with entries being expression levels. Ideally these data are processed using a uniform set of procedures. Second, the samples need to be described: what tissues, conditions etc. they were derived from, and what role they play in the study (e.g. control or treated samples). Third, for microarrays, the probes need to be mapped to genes (there is an analogous step for RNA-seq data), and this also should be done in a consistent manner.

The information provided by GEO is only a starting point to meeting these needs. Expression levels can be obtained from GEO for many datasets, but they are provided by submitters and thus are not processed in a consistent manner. Samples are described in GEO using free-text with limited structure provided by key-value pairings, but the descriptions are written by data submitters and are generally not adequate for our purposes, especially in the lack of harmonization that the use of formal ontologies affords. Mappings of microarray probes to genes are generally provided by data submitters (i.e., microarray manufacturers) and are often based on provided GenBank IDs for mRNAs. GenBank IDs rarely correspond to the actual sequences on the microarray, which are generally short oligonucleotides or (for some older platforms) PCR products.

To address these issues, Gemma implements a comprehensive data processing pipeline (Figure 1). Gemma natively handles data from GEO, but can take data from other sources formatted as a simple tab-delimited text file. In the following subsections describing our procedures, we do not give full details of every step and parameter, and we omit many software implementation details. Instead we attempt to document key steps and highlight aspects that are unique to Gemma. Additional details are available within the Gemma codebase (https://github.com/PavlidisLab/Gemma/).

**Figure 1:**
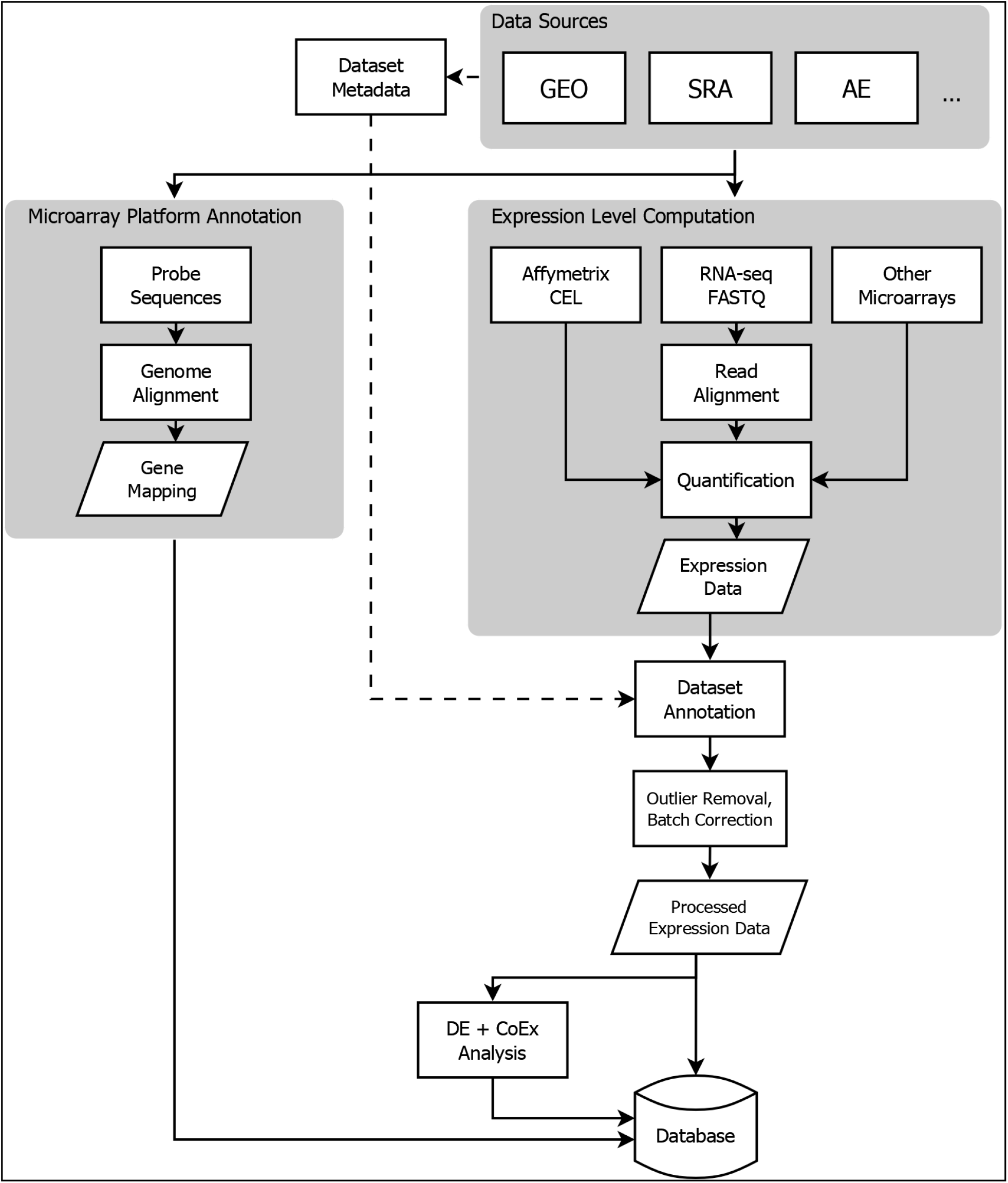
Flowchart showing the flow of data from NCBI Gene Expression Omnibus (GEO) into the Gemma database. Three classes of information are processed: platform metadata, gene expression data and metadata. While microarray data reside on GEO, raw RNA-sequencing data is obtained from NCBI Sequence Read Archive (SRA). Other abbreviations: ArrayExpress (AE), Differential Expression (DE), and Co-expression (CoEx).

#### Dataset selection

For Gemma, our goal is not to capture all available datasets, in part because of resource limitations. Initially, we accumulated data on any domain of biology for seven taxa of interest (*Homo sapiens*, *Mus musculus*, *Rattus norvegicus*, *Danio rerio*, *Drosophila melanogaster*, *Caenorhabditis elegans* and *Saccharomyces cerevisiae*). More recently, we have specifically increased our coverage of studies relating to the nervous system, and on datasets studying human, mouse, or rat. We de-prioritize datasets that have small sample sizes (i.e., less than 10 samples in total) as these are less suitable for the downstream analyses implemented in Gemma. We also prefer studies that have biological replication and a clear experimental design affording specific comparisons between contrasting conditions. That said, we do accommodate user requests for loading datasets that do not meet these criteria. A notable current omission is single-cell RNA sequencing (scRNA-seq) datasets, because most of the available datasets to date have minimal (if any) biological replication, and are often surveys intended for cell type identification. Additionally, there is little consistency in how single-cell datasets are represented in GEO, with some datasets having GSM identifiers for each individual cell, while others for each processed pool of cells. Anticipating that future scRNA-seq studies will eventually shift to designs with replicates, incorporating single-cell data into Gemma is a key area of future work.

#### Platform selection

Gemma can handle a wide range of microarray platforms including Affymetrix GeneChips, Agilent spotted arrays, Illumina BeadArrays and many two-color platforms, as well as shortread RNA-sequencing data such as Illumina sequencing-by-synthesis. Gemma does not yet have facilities to handle long-read sequencing data (e.g. PacBio or Oxford Nanopore), but since these are less commonly used and less suitable for quantification of expression levels, implementing support for these technologies have not been a priority. Gemma can also handle microRNA (miRNA) microarray platforms. Finally, we no longer routinely add data from two-color microarrays, partly because they are no longer in much active use. They also tend to have poor gene coverage and/or do not provide sufficient metadata to permit probe sequence analysis (see “Processing platform information”), as well as having other complications such as dye-swap or “loop” designs that hinder automated analysis.

#### Auditing and curation assistance tools

Gemma has a robust security model that permits control of ownership and visibility of underlying entities. All datasets when first loaded into Gemma are marked “private” and can only be viewed by curators and administrators. This allows us to curate and preprocess the data before making it available publicly. Gemma also has an audit trail system, in which updates and key events are recorded during processing. Curators can attach notes to datasets, mark datasets as needing curation, or flag datasets as unusable (we support most of these features for platforms as well). Platforms or datasets that are confirmed as unusable are blacklisted, to avoid inadvertently reloading them in the future. Collectively, these tools enable curators and administrators to easily assess the curation state of a dataset, track down the source of errors, and coordinate curation efforts.

### Processing of platform information

A key step in Gemma data processing is linking expression data to genes. This is especially important for microarray platforms, where the probe sequences on the arrays were often designed prior to the availability of high-quality reference genome sequences and annotations. We decouple the expression data from the underlying microarray platform, enabling us to update the relationships between probes and genes with minimal disruption.

We apply different probe-to-gene mapping strategies depending on the type of platform (e.g. Affymetrix probe sets, single oligonucleotide probes, and cDNA probes); this is done internally by adopting a previously described protocol (20) with some minor modifications. We first have to obtain the actual nucleotide sequences of the probes. These are often not available from GEO, thus we acquire them either from the manufacturers’ websites or by directly contacting them. Despite these efforts, for some platforms, the probe sequences remain unavailable. We blacklist these platforms as unusable because there is no way to confirm that the probe corresponds to the gene claimed. As these platforms tend to be used for very few datasets (often just one), the overall impact of this blacklisting is minor.

For platforms with probe sequences, the processing proceeds as follows. For Affymetrix platforms specifically, we perform an additional step where we collapse the Affymetrix probe sets (consisting of multiple 25-bp oligonucleotides each) to a single representative sequence, resolving overlaps, as described previously (20). Next, for all platforms, we then align the probe sequences against the appropriate reference genome with BLAT (21). Whenever a new genome assembly is available, the probes are realigned. The reference genome versions we currently use for human, mouse and rat are hg38, mm10 and rn6 respectively. Alignments are then evaluated and filtered for probes yielding specific alignments. Finally, probes are mapped to transcripts using genome annotations from UCSC GoldenPath (22). These mappings are also refreshed periodically to reflect updates to the UCSC annotations. Gene mapping details and links to visualization of alignments in the UCSC Genome Browser are provided through the platform’s page in Gemma, along with downloads of the annotations in a tab-delimited format. Overall, this procedure ensures (to the best of our ability) that probe annotations reflect the actual physical probe on the array, and are annotated in a consistent manner.

For RNA-sequencing data, a different strategy is employed while still accommodating it in the same general framework. For each taxon, we define a “pseudo-platform”, where the entire platform’s elements are the set of known genes recorded in the reference genome annotations (e.g. https://tinyurl.com/Gemma-HumanNCBI). The output of our RNA-sequencing data processing pipeline (see “Processing of expression data” below) can then be linked to these “generic” platforms based on NCBI gene IDs.

### Processing of expression data

#### Loading and pre-processing

First, basic dataset metadata (e.g. dataset title and description, number of samples, sample annotations etc.) are parsed and loaded from GEO SOFT files (Gemma can also load data from other sources *ad hoc* using a simple tab-delimited file). This is generally based on a GEO Series (GSE ID), and the system automatically retrieves any associated “GEO DataSets” (GDS IDs) if available. Unlike GEO Series, DataSets are curated by GEO, and provide additional metadata including basic information on the experimental design; they are however, only available for < 5% of GEO Series. Parsing of these files and mapping them to the Gemma data model results in an initial representation of the dataset in Gemma.

Next, expression level information is acquired. The default method extracts the expression levels included in the SOFT file at initial loading, if present. For datasets using two-color microarray platforms, we attempt to extract the data for both channels separately (including background levels) for assessing spot quality; spots which have low signal in both channels are flagged as missing data.

For Affymetrix microarrays, we reprocess the data from CEL files (when available) using the Affymetrix Analysis Power Tools provided by the manufacturer (https://tinyurl.com/Affy-APT; specifically “apt-probeset-summarize” with the RMA algorithm (23)). This data replaces that which was obtained from the SOFT file. We always use the manufacturer-provided probe groupings (“CDF” files); thus “custom CDF” analyzed datasets in GEO are reanalyzed using the standard CDF for the platform. This further enhances consistency compared to the user-submitted processed data in GEO.

For RNA-sequencing data, a separate pipeline (https://github.com/PavlidisLab/rnaseq-pipeline) is used to download FASTQ files from the Sequence Read Archive (SRA) or EMBL-EBI European Nucleotide Archive (ENA) (24,25), read adapters trimmed with Cutadapt (26), reads aligned with STAR (27), followed by quantification using RSEM (28). This also results in metadata on alignment statistics. Genelevel quantification data are then loaded into Gemma’s database.

After the data is loaded into Gemma (through either of the above streams), the resulting expression data is processed through a common post-processing pipeline, including steps described below. The expression data is log_2_-transformed (if not already) and then quantile normalized.

#### Batch correction

Where possible, the data are batch corrected, and is performed only after the curation of the experimental design. Importantly, batch information is almost never explicitly provided as sample annotations by the submitters in GEO. In Gemma, batches are defined automatically using information in the raw data files. For microarrays, date stamps can usually be found in CEL files for Affymetrix datasets; and in GenePix output files for Agilent (and other spotted platforms) datasets. Occasionally, batch information is instead obtained manually from supplementary information provided by the data submitter or the associated publication. Through the date stamps, batches are defined using simple heuristics (essentially a one-dimension clustering): samples that are processed closely in time are considered a separate batch if a larger time gap occurred before processing of the next sample. In practice this almost always yields a batch factor that is correlated with one of the first three principal components of the expression level data. For RNA-seq datasets, no relevant dates are associated with the records in GEO or in SRA. Instead, we attempt to use information from the FASTQ headers, which for Illumina platforms often contain information on the “device”, “run”, “flow cell” and/or “lane”. Any batch information obtained is stored as a factor in the experimental design. Despite these efforts, for many datasets we are unable to obtain batch information.

Batch correction is conducted using an in-house implementation of the ComBat algorithm (29), enhanced to automatically select covariates to include from the experimental design based on their loadings in the principal components of the data. Batch correction is not performed when batches are confounded with experimental design factors, or when no substantial batch effect is detected. Such cases are assigned special flags in the system that can be used during downstream analysis.

#### Diagnostics

A number of diagnostics are computed and stored for each dataset, such as principal component analysis (PCA) of the expression levels, the relationship between the principal components and curated experimental factors, the mean-variance relationship of expression levels (which is especially important for RNA-seq data (30)), and sample-sample expression level correlations. These are used for manual and automated QC processes, such as outlier detection, described next.

#### Outlier sample detection

A common data quality problem in expression datasets is the presence of one or more samples which deviate markedly (in non-biologically-relevant ways) from the properties of other samples. Gemma flags potential outlier samples automatically, which are then manually reviewed. Samples confirmed to be outliers are not permanently removed from the system: we represent them as missing data instead. This ensures transparency of the process, and also affords us the possibility to reverse the decision easily. Our outlier detection algorithm uses the sample-sample expression level correlation matrix, the idea being that samples which have low correlations with all other samples are potentially outliers. Omitting some details, the algorithm is as follows: the correlation matrix is first adjusted by regressing out the effect of major experimental factors such as tissue type, so that identifiable biological groups of samples are not treated as outliers. Samples are considered potential outliers if their adjusted median correlation is outside the inter-quartile range of correlations of the sample with the closest median correlation to them. All outliers called by the algorithm are reviewed and approved by a curator. With outlier removal and batch correction, the data is now finalized for downstream analyses.

### Multi-platform, multi-species, and overlapping datasets; GEO SuperSeries and SubSeries

Datasets in GEO are very diverse in how they are represented. Gemma is designed to automatically handle many complexities that arise; in some cases, a curator needs to review the dataset and take an appropriate action.

#### Overlaps

When a dataset’s SOFT file is downloaded from GEO and parsed in preparation for loading, Gemma checks to see if any of the samples are already included in another dataset in Gemma, based on the “GEO Sample” ID (GSM IDs). Any duplicated samples are removed and noted in the dataset description (this does not catch cases where the same data is submitted to GEO twice, yielding different GSM IDs, though we have caught such cases by manual curation). Occasionally, this results in a suboptimal representation of either or both datasets, e.g. it can result in the removal of all control samples from a dataset. To resolve this issue, we determine which of the overlapping datasets results in the best use of the samples, and then reload the “better” dataset to have a usable design.

#### Multi-species

Datasets that include samples from multiple species are detected and split apart so that a separate Gemma ExpressionExperiment is created for the samples from each taxon (e.g. GSE23579 (31) consists of both human and mouse samples, and are split into two Gemma datasets: GSE23579.1 [https://tinyurl.com/Gemma-GSE23579-1] for human, and GSE23579.2 [https://tinyurl.com/Gemma-GSE23579-2] for mouse). Samples from taxa that are not supported by Gemma are rejected.

#### Multi-platform

A common occurrence in GEO is that a dataset uses more than one platform. For datasets that include non-expression profiling data, only the transcriptome samples are retained. We then apply procedures to resolve the use of multiple transcriptome profiling platforms in the dataset. First, if the platforms are considered incompatible, the dataset will be split by the platforms, similar to how we handle the multi-species case. An example would be the use of a two-color spotted array platform and an Affymetrix platform in the same dataset. If the platforms are compatible (e.g. all Affymetrix platforms, or multiple generations of Illumina short-read sequencing platforms), then the data from the two platforms are combined. There are two major sub-cases. In the first case, each sample was run on a different platform, e.g. some samples were run on the HG-U95A platform and others on HG-U95Av2. These two platforms are very similar, so a combined dataset can be constructed where common probe sets have data for all the samples. The second case arises with multi-part microarray platforms, such as HG-U133A and HG-U133B, where samples were run on both platforms. In this case a combined dataset is constructed by “stacking” the data for the two platforms. This is however not straightforward: using the same example platforms, deciding which HG-U133A sample matches the corresponding HG-U133B sample is nontrivial, because GEO does not model the concept of an RNA sample, and thus there is nothing formally linking the two samples together. Gemma has tools to assist matching samples across platforms automatically (through a heuristic based on sample names and metadata), followed by manual review; alternatively, the matching step can also be performed entirely manually. Once resolved, such datasets are represented in Gemma as being conducted on a single merged platform that encompasses all of the platform elements.

#### SuperSeries and SubSeries

GEO has the concept of SubSeries, which are grouped together to make up a SuperSeries. However, the semantics of this relationship are not consistent. Sometimes a SuperSeries is made up of related but separate datasets, and this is probably how it is intended to be used; but in other cases, GEO submitters use the concept to represent an experimental design, such as putting all control samples in one SubSeries and all treated samples in another SubSeries. Because of these kinds of issues, we manually review SubSeries and SuperSeries datasets to decide whether they should be imported into Gemma at the SuperSeries level or as separate SubSeries datasets.

#### Other instances of dataset-splitting

As described above, we often divide GEO GSE records into more than one Gemma dataset based on species, platform, or SubSeries. However, there are other cases where it is desirable to split datasets. One example is when there is a batch-confound with the experimental design, and splitting the dataset by the “batch” factor would yield multiple usable sets of samples. A factor often found confounded with the batch factor is tissue (“organism part”). By splitting such datasets, we sacrifice the ability to do tissue comparisons, but salvage other aspects of the study. Another situation that arises is when the GEO submitter should have represented their study as multiple GEO series (perhaps SubSeries), or when an otherwise “clean” experimental design is disrupted by the inclusion of an “extraneous” sample, such as a technical control from a non-relevant cell line. The ability to split datasets *ad hoc* is a relatively new feature in Gemma, and has not yet been applied to many studies.

### Gene Expression Experiment Quality (GEEQ) Score

In the course of curating datasets with wildly varying properties, we decided it would be beneficial to provide a summary of various considerations to give a “gestalt” sense of data quality, which we call GEEQ scores. The GEEQ score was developed to reflect the qualities of good data in a topic-agnostic manner. The quality score is meant to help answer the question “How well can I trust the results of an analysis of these data?” The features that are used to compute the score and their weightings are described here (https://pavlidislab.github.io/Gemma/geeq.html). Some of the properties we take into account include the number of replicates for each condition, severity of batch effects (unless corrected), and median sample-sample correlation. Prior experience has shown that datasets that have issues in these categories tend to give results that are noisy and less reproducible; we assign a lower GEEQ score to reflect that. Based on extensive experience, we calibrated the GEEQ scores such that values in the lowest and highest quintiles reflect the extremes of observed data quality. These scores are merely a rough guide to assist users in identifying datasets that might be especially suitable or problematic.

### Curation of dataset metadata

After loading and basic preprocessing has been done to the extent possible via automation, every dataset is manually curated. Our manual curation is largely performed by trained undergraduate research assistants who follow a written set of guidelines. Here we describe the curation of dataset metadata, which encompasses annotating both the experimental design and the “topics” of the dataset. Manual quality control checks (i.e. batch correction and outlier removal) as described earlier are performed after metadata curation.

The task of curating a large number of datasets consistently by different curators is a challenge in balancing different priorities. The process itself should be fast, while capturing sufficient experimental information for dataset retrieval and downstream analysis (i.e. differential expression), and maintaining enough consistency to facilitate cross-dataset comparisons. We developed a set of curation guidelines that accommodate many types of commonly-encountered dataset topics and experimental designs. The guidelines provides instructions for correctly mapping a dataset’s experimental design into the appropriate representation in Gemma (including the level of detail to be used and standardizing the choice of ontology terms used during annotation), and guidance on prioritizing dataset topics for annotation. New rules are introduced only when absolutely necessary, to avoid the guidelines from becoming too unwieldy.

Over the next few subsections, we will describe the metadata curation guidelines. First, we describe the rationale for using ontologies and our approach of doing so. Second, we describe the specific details for design- and dataset-level metadata curation respectively. Third, we describe how the dataset search system is supported by our use of ontologies.

#### Use of ontologies

For both tasks of annotating and searching datasets, it is advantageous to use formal ontologies. Ontologies are created to capture various concepts in the form of “terms”, and elucidate the relationships between those concepts; both of which are performed in a standardized manner. Cross-ontology relationships are also captured through the use of “concept imports”. By using ontologies during metadata annotation, we have more assurance that the design and topics are captured in a standardized manner, and provides potential for interoperability with external resources that utilize the same ontologies. Similarly, we can harness the strengths of ontologies during searches, especially through inference of the ontology’s graph structure (see “Dataset search”). Our use of ontologies in Gemma is semantically simplistic and is best represented as key-value pairs: an entity is annotated with a “Category” and a “Value”. For example, if a sample was treated with vincristine it would be annotated with Category = “treatment” [EFO_0000727] and Value = “vincristine” [CHEBI_28445]. Multiple annotations of a single entity are treated as a “bag-of-pairs”, with no explicit semantics linking them, e.g. 10μM of vincristine has the Value = {“vincristine” [CHEBI_28445], “10 uM”}. This design decision favors simplicity of implementation and speed of curation at some cost of power for inference. Next, we will discuss the standards used for the “Category” and “Value” pairs separately.

#### Category standardization

As shown in the previous example, we include “Categories” when annotating Gemma entities. This is done for two key reasons: by assigning a category to the terms’ “Value”, we provide additional context to those terms, and we enable easier grouping of entities. To illustrate the point: if a sample is annotated with the gene symbol *IGF1*, assigning the category “genotype” indicates that there is a genotypic modification of the gene *IGF1* in the sample’s cells; whereas assigning the category “treatment” indicates that external IGF1 proteins were introduced into the sample’s culture media. In terms of grouping entities, when “categories” are used as factors, it enables us to group samples by the respective factors’ levels and perform differential expression analysis. Additionally, categories can be used as filters when searching for annotations in Gemma. Thus, it is essential that the use of categories is standardized. In Table 1, we list the mapping of categories to their appropriate category ontology term (mostly derived from Experimental Factor Ontology (EFO) with some exceptions) and examples of concepts in which the category is applied.

**Table 1:** Table of concept categories, their assigned category ontology term and examples of concepts for use during Gemma curation. Category ontology terms are mostly derived from terms in the Experimental Factor Ontology (EFO), with some exceptions derived from the Phenotype and Trait Ontology (PATO) and the Cell Line Ontology (CLO).

| Concept Category | Category Ontology Term | Concept Examples |
| --- | --- | --- |
| Genetic Manipulation | Genotype [EFO_0000513] | Gene knock-outs/-downs, over-expression. |
| Chemical/Physical Treatment | Treatment [EFO_0000727] | Drugs, burns. |
| Tissue/Organ | Organism Part [EFO_0000635] | Liver, brain. |
| Cell Type | Cell Type [EFO_0000324] | Hepatocytes, astrocytes. |
| Disease | Disease [EFO_0000408] | Breast cancer, Parkinson’s disease. |
| Disease Stage | Disease Stage [EFO_0000410] | Metastatic cancer. |
| Mouse/Rat Strain | Strain [EFO_0005135] | C57BL/6, Long-Evans. |
| Sex | Biological Sex [PATO_0000047] | Male, female. |
| Cell Line | Cell Line [CLO_0000031] | HeLa, MCF7. |
| Sampling Time | Timepoint [EFO_0000724] | 5 hours, 10 minutes. |
| Developmental Stage | Developmental Stage<br>[EFO_0000399] | Embryonic Stage Day 5 (E5),<br>Post-Natal Day 2 (P2). |
| Age | Age [EFO_0000246] | 25 years, 70 years. |

#### Ontology selection

Gemma currently supports 12 different ontologies (Table 2). There were multiple considerations in selecting ontologies that would be supported by the system. Curators must be able to search the ontologies and identify the concepts they need to use, and having too many ontologies makes this more difficult. Having too many ontologies also impacts search performance (speed and precision). For example, we decided to support the Uberon ontology (32), in lieu of the extensively overlapping Foundational Model of Anatomy (FMA) ontology (33). Despite choosing ontologies judiciously, there remain duplicated concepts across ontologies (e.g., the HeLa cell line exist in both the Experimental Factor Ontology [EFO_0001185] and Cell Line Ontology [CLO_0003684]). Our curation standards help enforce consistency. Thus for example, cell lines are annotated with CLO terms as far possible. In Table 2, we list the general groupings of concepts, and the recommended source ontology in which the terms (to represent those concepts) are to be used during curation.

**Table 2:**
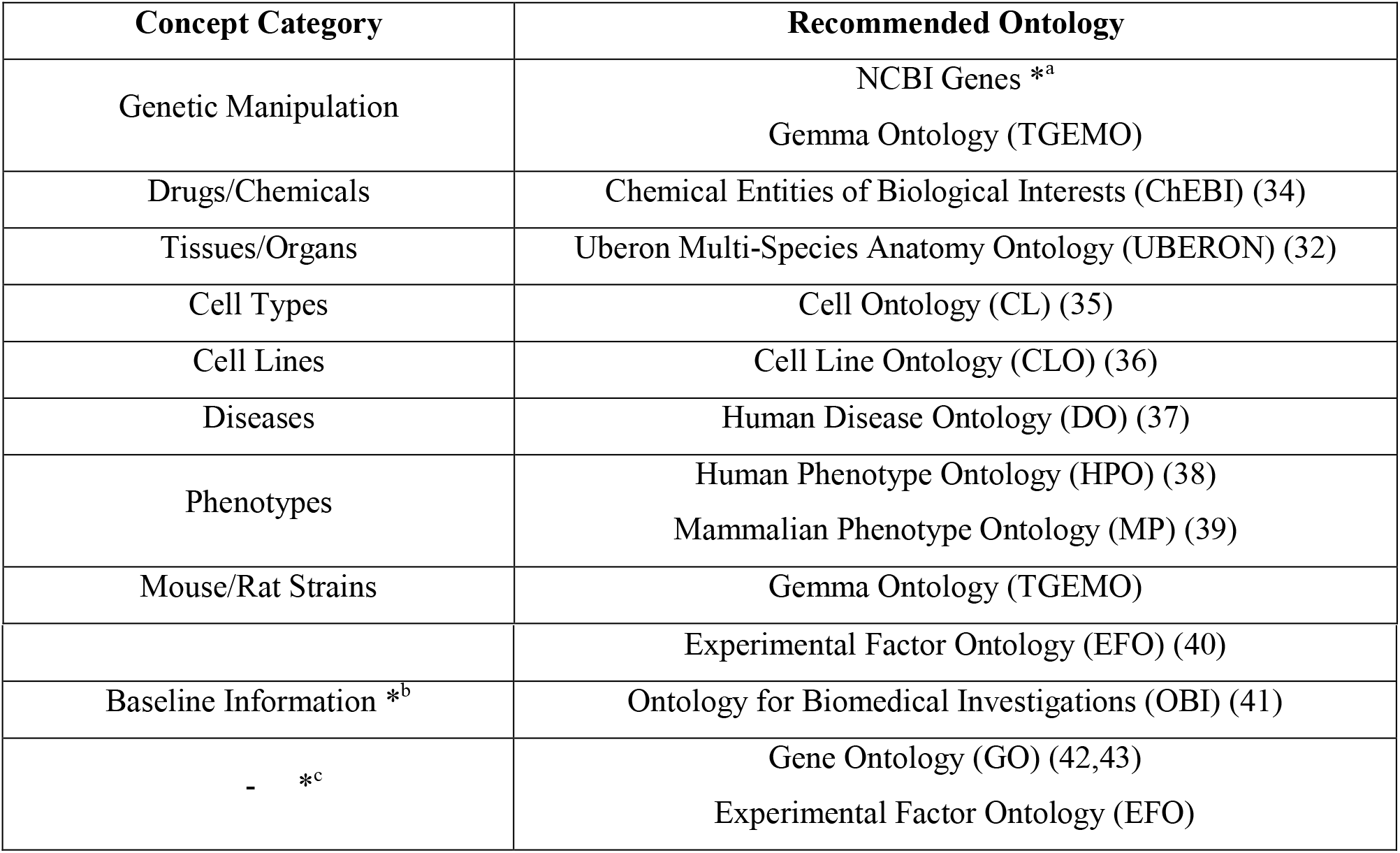
Table of concept categories and their recommended source ontologies for use during Gemma curation. Strictly speaking, there is no formal NCBI Gene ontology: we generated a “pseudo-ontology”from the gene accession IDs to represent the genes of the seven taxa supported by Gemma (*a). “Baseline” concepts are used exclusively for design-level annotation, while the others are used for both design- and dataset-level annotation (*b). For concepts that are infrequently occuring or could not be grouped, we usually default to using terms in Gene Ontology and Experimental Factor Ontology whenever possible (*c).

| Concept Category | Recommended Ontology |
| --- | --- |
| Genetic Manipulation | NCBI Genes <sup>*a</sup><br>Gemma Ontology (TGEMO) |
| Drugs/Chemicals | Chemical Entities of Biological Interests (ChEBI) (34) |
| Tissues/Organs | Uberon Multi-Species Anatomy Ontology (UBERON) (32) |
| Cell Types | Cell Ontology (CL) (35) |
| Cell Lines | Cell Line Ontology (CLO) (36) |
| Diseases | Human Disease Ontology (DO) (37) |
| Phenotypes | Human Phenotype Ontology (HPO) (38)<br>Mammalian Phenotype Ontology (MP) (39) |
| Mouse/Rat Strains | Gemma Ontology (TGEMO) |
|  | Experimental Factor Ontology (EFO) (40) |
| Baseline Information *<sup>b</sup> | Ontology for Biomedical Investigations (OBI) (41) |
| - *<sup>c</sup> | Gene Ontology (GO) (42,43)<br>Experimental Factor Ontology (EFO) |

During the curation process, we found that a number of concepts are not represented in the ontologies we use. We created The Gemma Ontology (TGEMO; https://github.com/PavlidisLab/TGEMO) to capture these terms, with the intent of replacing them with the appropriate ontology terms in the future. Currently, TGEMO contains 101 ontology terms, and implements concepts for genetic modifications (e.g. Knockdowns), some mouse/rat strains (e.g. DB/2) and some terms for our internal usage.

#### Free-text annotation standards

While the coverage of concepts by the ontologies we use is fairly extensive, there still are concepts for which we resort to free-text. These are predominantly numerical measures, such as developmental stages of an organism (e.g. E3, P4, etc.), sampling time intervals (e.g. 3 hours) and dosages (e.g. 3 μg/ml, 2 mM/ml). We established style standards for free-text annotations to ensure consistent and unambiguous annotation (e.g. usage of SI units and abbreviations, appropriate spacing, official nomenclature guidelines etc.)

Separately, while our design decision of representing characteristics as “bag-of-pairs” is adequate for the majority of use cases; this is less so for more complex situations, such as some genetic experiments. For example, in GSE1463 (https://tinyurl.com/Gemma-GSE1463, (44)), there is a group of samples in which both the *Dmd* and *Utrn* genes were simultaneously knocked out. Without the free-text annotation, it might be possible to infer from the ontology terms used that both genes were knocked out; this is made more explicit and unambiguous with the free-text annotation. Another example is GSE52022 (https://tinyurl.com/Gemma-GSE52022, (45)), where mutated human *APP* and *PSEN1* genes were introduced and expressed in transgenic mouse models. With the free-text annotation, it is clear which mutations belonged to which genes, and the entire mouse genotype can be easily read.

#### In-built concept curation support

Gemma has built-in search support for the use of ontologies during curation. When a curator is inputting the text values of an ontology term to express a concept (e.g., a drug), the input box automatically searches and provides a list of recommended matching terms, along with an indication as to whether (and how often) the term has been used previously in the system. Frequently-used terms are favoured. When no suitable term is identified, the curator can choose to use free-text to annotate the concept by following formatting guidelines of common cases. This also promotes the reuse of previously used free-text strings, which are also presented in the concept recommendation results (albeit with a lower ranking).

#### Experimental design curation

In Gemma, the concept of an “experimental design” refers to the characteristics of a dataset’s samples, in which groups can be formed for downstream analysis of the data, such as differential expression. These characteristics are used as input in our statistical models (see “Downstream analysis”), and also in dataset searches. Experimental designs are modeled as follows. A dataset can have one or more ExperimentalFactors that describe a feature (or factor) that varies across samples; while the different levels of the feature are called FactorValues. For example, the dataset GSE2198 (https://tinyurl.com/Gemma-GSE2198; (46)) has three factors: “organism part” for different tissues, “genotype” for different genotypic states, and “block” for sample batches. With the exception of “block” which has four levels (i.e. four sample batches); both “organism part” and “genotype” have three levels each. Gemma supports both categorical and continuous factors, though nearly all factors are categorical.

Most datasets, when first imported into Gemma, do not have any information on the experimental design. Exceptions to this are datasets (specifically GEO Series “GSE”) that have an associated GEO DataSet “GDS”, which have been manually curated by NCBI GEO staff. For such datasets, we automatically initialize an experimental design layout based on that in the GDS. In either case, we manually curate the experimental design to add ontology terms and to enforce uniformity in how factors and levels are described.

Considering the diversity in experimental design of the datasets submitted to GEO, we elect to prioritize key experimental factors during curation. The rationale is that while we could invest more time to exhaustively curate every single potential factor within a dataset, it would be a large effort with diminishing returns. Instead, we prioritize factors that are highly recurrent across many studies which would give us the greatest value. These factors include: “genotype” for genetic manipulations; “treatment” for both chemical and physical treatments; “disease” for diseases and disorders; and “disease staging” for the varying stages or severity of disease. For time-course datasets, factors such as “timepoint” for sampling time points; “developmental stage”; and “age” for patient ages are also prioritized. Other popular factors that are taken into account include “organism part” for tissues and organs; “cell types”; “cell lines”; “strain” for mouse or rat strains”; and “biological sex”.

Another aspect of design curation is the identification of the “baseline condition” of the study, which is important for differential expression analysis. Assignment of the baseline condition is not performed for all factors, as there might not be any to begin with (e.g. “organism part”, “biological sex”). For factors that do have a baseline condition (e.g. “genotype”, “treatment”, and “timepoint”), the condition is annotated with specific ontology terms such as “reference substance role” etc. (see “Downstream analysis”). For more complex datasets which involves positive and negative controls, we use the “control” term to annotate what we perceive as the “true” baseline condition of the dataset. For example, in dataset GSE40463 (https://tinyurl.com/Gemma-GSE40463; (47)), for one of the “genotype” factors, there were four levels: over-expression of *Tbx21*, over-expression of *Gata3*, presence of the empty transgene vector, and a negative control without the transgene vector. Since the first two levels (i.e. transgene overexpression) rely on the presence of a transgene vector, we consider the samples with the empty transgene vector to be the baseline condition.

#### Dataset topic tags

Dataset-level annotations are used to capture information that is often not explicitly part of the experimental design and not otherwise captured. An example would be a tissue or disease state that is constant across all of the samples in a dataset, but also relevant topics that might not be otherwise explicit. These tags are used to help ensure that searches for a relevant concept would retrieve the dataset. For example, GSE36051 (https://tinyurl.com/Gemma-GSE36051), a study of breast tumor cells (48), is annotated with Disease Ontology terms “breast cancer” [DOID_1612]. Similarly, relevant disorder terms are tagged on datasets using animal models, such as GSE14499 (https://tinyurl.com/Gemma-GSE14499), a dataset concerning a mouse model for Alzheimer’s disease (49). In this case, the Disease Ontology term “Alzheimer’s disease” [DOID_10652] is used because even though the study is not directly studying Alzheimer’s disease, it would be reasonable for users searching for Alzheimer’s disease to see this dataset in the results. In keeping with this philosophy and to avoid unnecessary curation, terms that are used in annotating the experimental design are not repeated in the topic tags, and vice versa, since both are available to the search engine. We ensure that certain types of information are always captured including tissue of origin, cell type, cell line, mouse/rat strains and biological sex.

#### Dataset search

An important use case for Gemma is searching for datasets based on keywords. The system we have developed performs search based on both full-text indexing as well as ontology inference. In text searches, we search all dataset-associated text fields: title, description, sample names, etc.; while ontology searches are performed in two different ways: free-text searches are used to retrieve ontology terms based on an index of the ontology-associated text (via Apache Jena’s LARQ package; https://tinyurl.com/JENA-LARQ), whereas ontology URIs (Uniform Resource Identifier) can be used directly as search terms. In either case, retrieved ontology terms are expanded to include all child terms via “part-of” and “is-a” relationships. This means that searching for the term “cerebral cortex” will not only retrieve datasets that contain that term either in free-text or in ontology term, but also datasets associated with the term “hippocampus” (in both free-text or ontology term), since “hippocampus” [UBERON_0001954] is a child term of “cerebral cortex” [UBERON_0000956].

### Downstream analysis

At this point in the processing pipeline, the data have been processed (normalized and batch corrected), the experimental design and other relevant metadata populated, and a final manual review has been performed.

Next, differential expression analysis is performed on the dataset based on the annotated experimental design. In cases where certain terms are used (e.g. “reference substance role” [OBI_0000025], “reference subject role” [OBI_0000220], “initial time point” [EFO_0004425], “wild type genotype” [EFO_0005168], “control” [EFO_0001461], etc.), Gemma automatically assigns these conditions as the baseline control group; in absence of a clear control condition, a baseline is arbitrarily selected. To perform the analysis, a generalized linear model is fit to the data for each platform element (probe/gene). For RNA-seq data, we use weighted regression, using an in-house implementation of the voom algorithm (30) to compute weights from the mean-variance relationship of the data. Contrasts of each condition are then compared to the selected baseline. In datasets where the “batch” factor is confounded with another factor, separate differential expression analyses are performed on subsets of the data; the subsets being determined by the levels of the confounding factor. This is reasonable for many cases we have observed (as mentioned earlier, in some cases, splitting the dataset is a better approach). One common situation is where different tissues (“organism part” [EFO_0000635]) were analyzed in batches.

The output of the differential expression analysis is then stored in the database. For each platform elements, this includes: At the factor-level, the main effect’s p-value and associated false discovery rate (q-value); at the contrast level, the test statistics (t-statistic); the log_2_ fold-change and its associated p-value. The results are available for download as tab-delimited files, and the top-most differentially expressed platform elements can be visualized on Gemma.

The other major type of analysis supported by Gemma is co-expression. This analysis has been performed on selected datasets that have sufficient number of samples (generally at least 20). As the co-expression component of Gemma is currently undergoing redesign, we omit a detailed description of the associated methods.

### Manuscript-specific code

Figure generation and numerical reporting was performed with code using Jython, Python, and R. Scripts and relevant data files are provided on Github (https://github.com/PavlidisLab/GemmaPaper-2020).

## Results

### Dataset and sample statistics

As of June 2020, Gemma provides 10811 curated gene expression datasets, with a total of 395,419 independent samples. These datasets span seven major taxa (*Homo sapiens,Mus musculus, Rattus norvegicus*, *Danio rerio*, *Drosophila melanogaster*, *Caenorhabditis elegans* and *Saccharomyces cerevisiae*). For most of this paper, we will focus our analyses and descriptions to the three primary taxa: human, mouse, and rat; each having 4593, 4933, and 894 datasets respectively (Figure 2A). Similarly, they account for most of the samples in Gemma, with human, mouse and rat totalling 237K, 123K, and 26K samples respectively (Figure 2B). All of these datasets were generated using either microarray or Illumina RNA-sequencing technologies, with the majority being microarray datasets (8429/10420, 81%; accounting for 342,635/385,397 or 89% of all samples). Almost all datasets (10351/10420, 99.3%) were obtained from NCBI GEO and are derived from 10288 unique GEO Series (i.e. GSE); with the balance coming from manual uploads of data from other sources (e.g. ArrayExpress). Putting the numbers in perspective, Gemma contains 14% of all expression profiling studies available on GEO for these three taxa. Neuronal-related datasets (largely excluding brain cancer) currently accounts for about 34% of all datasets within Gemma; as a point of comparison, 17% of Gemma datasets are cancer-related. Further description of the dataset topics is presented in a later section (“Dataset topics”). Aside from the publicly-available 10811 datasets, Gemma contains another 484 datasets that have been blacklisted and flagged for eventual deletion. At the individual dataset level, the number of samples varies greatly: for the bulk of datasets (90%, 9379/10420), the sample count ranges from 6 to 2158 samples, with a mean of 40.5 (Figure 2C). While the range appears large, only 6.5% (673/10420) of datasets have 100 or more samples each, and the distribution of sample number per dataset is similar across the three taxa of interest, with the human datasets tending to have marginally more samples than mouse and rat (Figure 2C).

**Figure 2:**
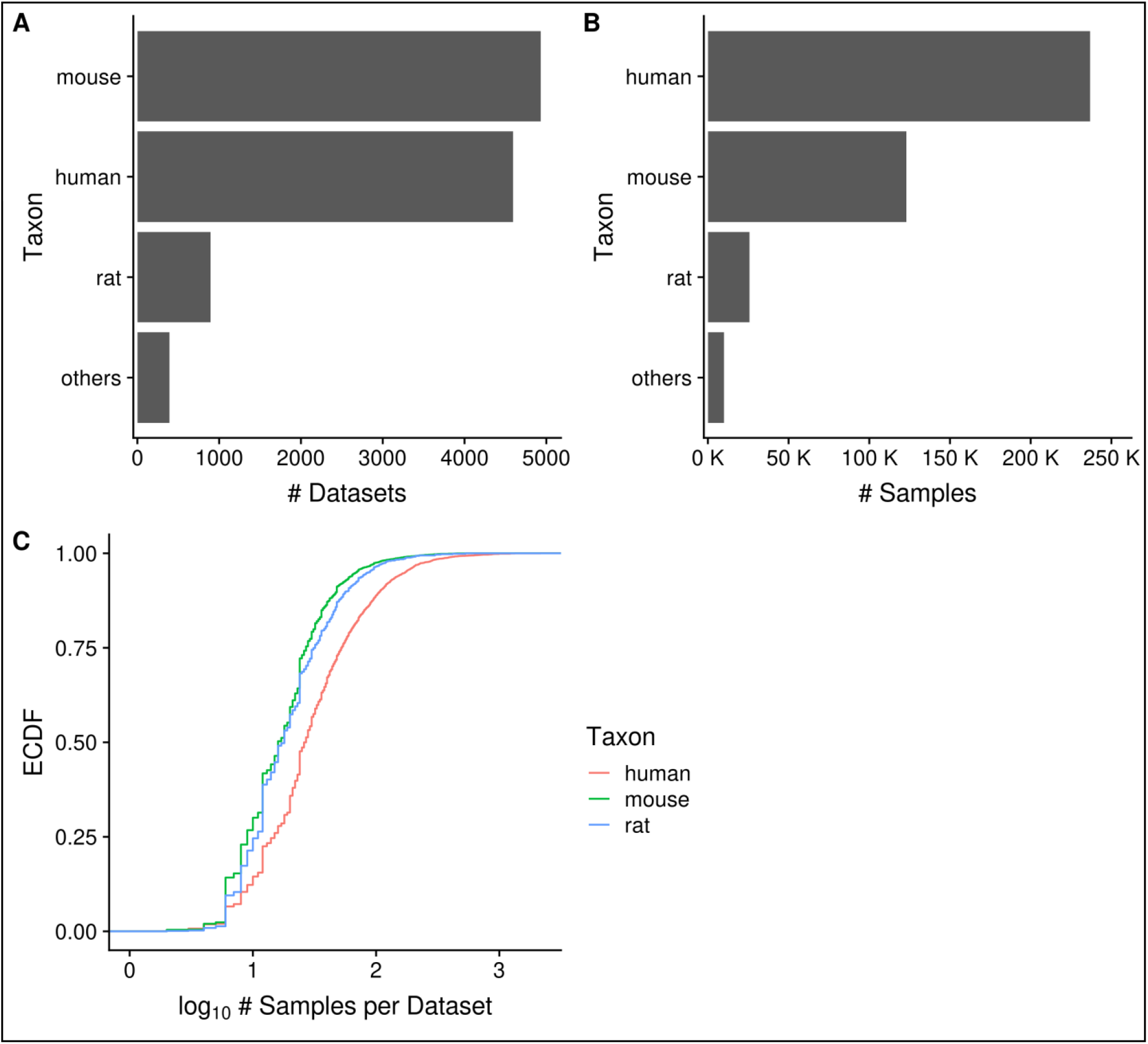
Horizontal bar charts displaying the number of datasets (A) and samples (B) grouped by taxon; Empirical cumulative distribution function (ECDF) representation of the number of datasets against the number of samples per dataset grouped by taxon (C). “Others” taxon group consists of *D. rerio, D. melanogaster, C. elegans* and *cerevisiae*. (N_dataset_ = 10420 in C).

### Dataset quality

Over the course of curating datasets, we noticed that studies vary widely in their data quality. We developed the Gene Expression Experiment Quality (GEEQ) scoring scheme as a means for quantifying the data quality (see Methods). These scores are publicly viewable (see User Interface), thus researchers could use these scores in determining which datasets of interest they could focus on (or omit) in their downstream analyses. We stress that these scores are only a rough guide, and a dataset’s quality score can sometimes be improved by curation.

Multiple dataset properties are used in determining the GEEQ score. Here, we present some additional data for three of the properties: median sample-sample correlation, presence of outliers in the dataset, and “batching” information. It is well established that a high correlation of gene expression profiles between replicate samples is a good indicator of data reproducibility (50,51). Using the same logic, while many datasets contain samples of different conditions (e.g. treatment-control comparisons), we expect to see strong agreement (i.e. high median sample-sample correlation) between biological replicates of the same condition, and in general after adjusting for known covariates, within a dataset. In Figure 3, we observe that the distribution of median correlations is centred at 0.91, with a mode of 0.99; the range of correlation spans from −0.7 to 1.0. This indicates that for most datasets (8512/10420, 82%), the within-dataset sample agreement is strong (ρ_median_ ⩾ 0.9). As high sample agreement is indicative of data quality, such datasets are given a higher GEEQ score. For the next property, 8% (825/10420) of datasets had at least one outlier sample flagged and manually confirmed during QC. The number of outliers removed ranges from 1 and 21 samples in those datasets; when converted to a ratio of outlier samples over the total number of samples in each dataset, the outlier removal ratio ranges from 0.002 and 0.36, with a mean ratio of 0.04. In relation to the GEEQ score, only the presence of predicted outliers negatively affects the GEEQ score: once these are reviewed (and potentially confirmed), they have no impact on the final score. Finally, 62% (6460/10420) of all datasets contain inferred batch information, of which 47% (3047/6460) were judged as not requiring batch correction, 19% (1231/6460) were batch corrected and the remainder (34%, 2182/6460) could not be corrected due to confounds between the batches and another experimental factor. It is concerning that almost 38% of all datasets lack batch information, which raises questions on whether the data can be fully trusted (and accordingly we downgrade these datasets with lower GEEQ scores); we found that this is prevalent in datasets using the Illumina BeadArray technology (1330/1356, 98%), as the submitted data files (both raw or processed) and metadata in GEO almost never contain any batch information or date stamps. We have observed that a fair number of batch-confounded datasets could be “rescued” by splitting the datasets on one of the factors that the original submitters did not intentionally control for (often times either by tissue or cell line). Rescuing of these datasets is a recent addition to our curation process that is still being put into effect.

**Figure 3:**
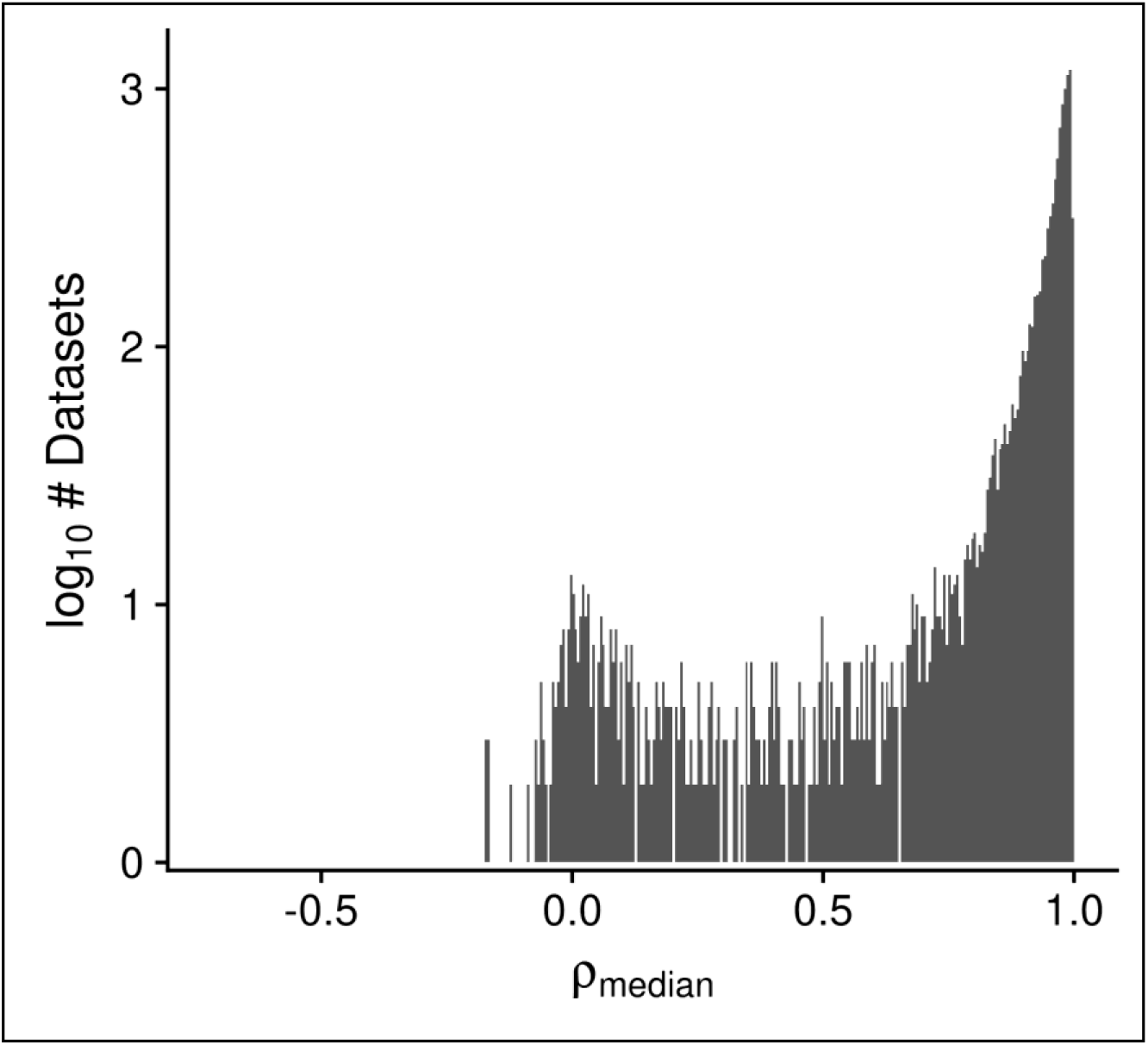
Histogram displaying the distribution of the median design-regressed sample-sample correlation per dataset, for all human, mouse and rata datasets. (N_dataset_ = 10420).

Separate from the GEEQ score framework, we explored two other dataset properties that may influence the dataset and curation quality: availability of raw expression data and association of a PubMed publication to the dataset. For the first, 64% (6686/10420) had raw expression data that was successfully reprocessed through our pipeline (70% of which were microarray platforms (4704/6686)). Since all of our RNA-sequencing data are reprocessed, this indicates that 56% (4704/8429) of microarray data had raw data that could be successfully reprocessed. For the second, 86% (8981/10420) had an association with a PubMed publication ID, indicating that for a vast majority of datasets, our curators will be able to better curate those datasets using additional information from the publications.

### Platforms

In Gemma, a total of 811 platforms have been loaded, though only 357 platforms are linked to datasets; the remaining 454 platforms are not used actively in Gemma for various reasons which we will discuss. Out of the 357 fully curated platforms, 92% (329/357) of them are used in human, mouse or rat (N = 150, 125 and 54 for human, mouse and rat respectively). All but seven are microarray platforms (the remainder being RNA-sequencing “pseudo-platforms”). Gemma also contains 21 “merged platforms” to accommodate datasets in which the same sample is analyzed on different (but closely related) platforms.

In GEO, RNA-sequencing data are linked to far numerous platforms due to the various iterations of Illumina short read systems; in Gemma however, these platforms (53/454 unused platforms) are collapsed into a single taxon-specific “pseudo-platform” that contains all known genes of the taxon (see Methods). Similarly, there are 88/454 alternative Affymetrix “platforms” that are listed separately in GEO despite using the same physical array, because they were processed using “custom CDFs”; datasets using these are remapped in Gemma to use the “official” manufacturer-designated probe set layout, except in cases where raw CEL files are not available. A further 160/454 unused platforms have no datasets currently linked with them and are retained for record keeping (many of these platforms are eventually merged). 34% (153/454) of the remaining platforms have been considered unusable for various reasons, such as microarray platforms lacking specific probe sequence information, and have been blacklisted. Much of the processing and harmonization of platforms in Gemma reduces the complexity of managing platform redundancies and improve platform comparability across technologies within each taxon.

The platforms with the most associated datasets for human, mouse and rat respectively are GPL570 (Affymetrix GeneChip Human Genome U133 Plus 2.0 Array), mouse RNA-Seq and GPL1355 (Affymetrix GeneChip Rat Genome 230 2.0 Array; N = 1335, 1325 and 263 respectively; Figure 4C). There are many platforms that have very few datasets run on them: 42% (137/329) of platforms are associated with only a single dataset (and this is nearly always representative of the platform’s usage frequency in GEO); 79% (259/329) of platforms are associated with ten or fewer datasets (Figure 4A). On the other hand, 7% (22/329) of platforms are associated with 100 or more datasets each, and taken together represent 78% (8079/10420) of all the datasets (human, mouse and rat) contained in Gemma. We also observe that microarray platforms that have more datasets tend to have a slightly better coverage of protein-coding genes (Figure 4B); the mean fraction for protein-coding gene coverage of microarray platforms with 100 or more datasets is 0.71 or 71% of known protein-coding genes, with a range from 0.33 to 0.94. In other words, 72% (6096/8429) of microarray-datasets (human, mouse and rat) contain data for on average 71% of known protein-coding genes.

**Figure 4:**
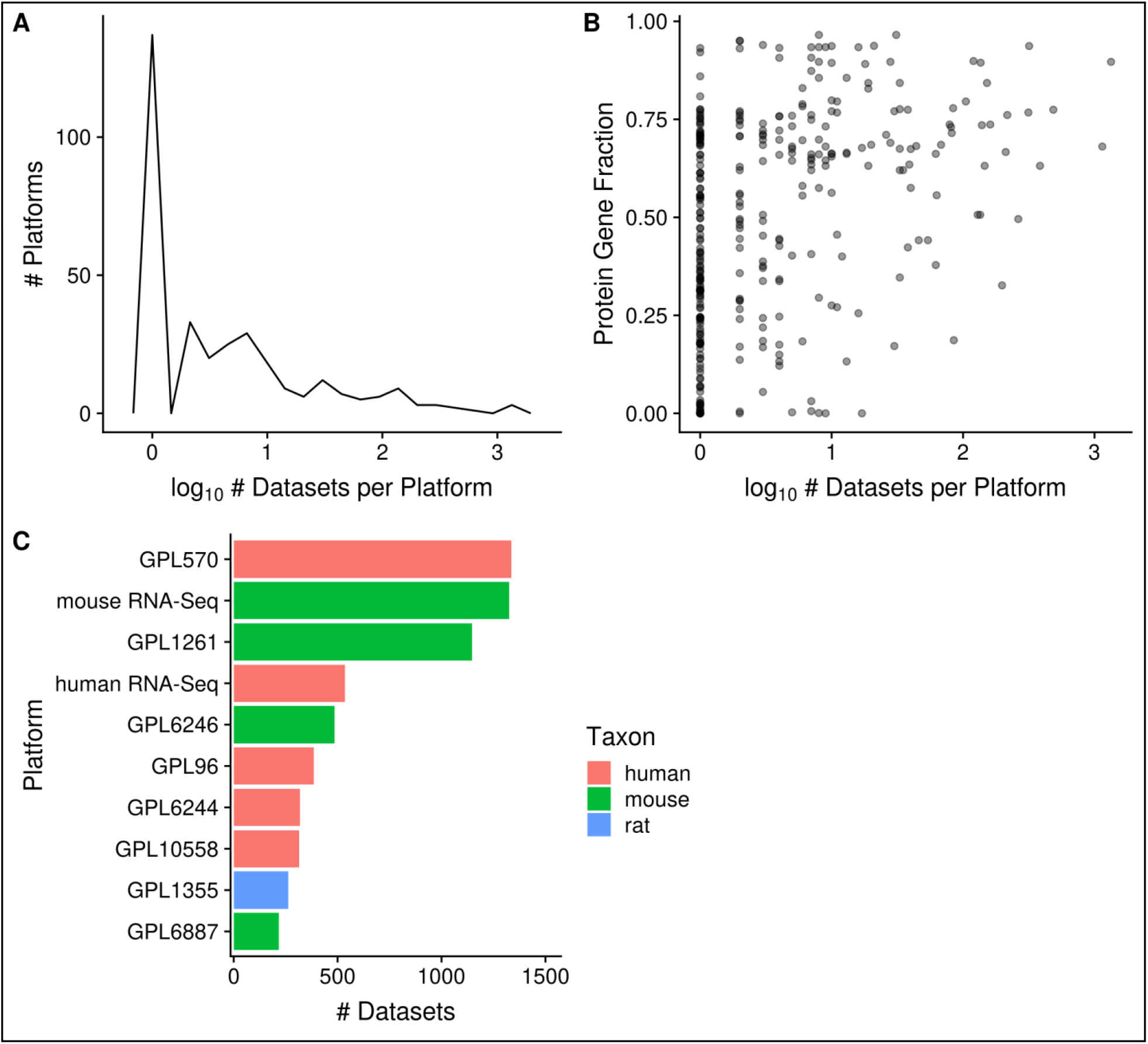
Distribution of platforms by the number of associated datasets for each platform (N = 357; A); Scatter-plot of the fraction of protein-coding gene coverage of each microarray platform with its number of associated datasets (N = 354; B); and bar chart of the top 10 platforms with the most associated datasets for human, mouse and rat (C). The names of the listed platforms (C): Affymetrix GeneChip Human Genome U133 Plus 2.0 Array (GPL570), Affymetrix GeneChip Mouse Genome 430 2.0 Array (GPL1261), Affymetrix Mouse Gene 1.0 ST Array (GPL6246), Affymetrix GeneChip Human Genome U133 Array Set HG-U133A (GPL96), Affymetrix Human Gene 1.0 ST Array (GPL6244), Illumina HumanHT-12 v4.0 Expression Beadchip (GPL10558), Affymetrix GeneChip Rat Genome 230 2.0 Array (GPL1355), and Illumina MouseWG-6 v2.0 Expression Beadchip (GPL6887).

### Dataset topics

Dataset topics are best understood as conceptual terms that are relevant to a particular dataset, and are displayed in a “bag-of-terms” model on Gemma. Across the curated datasets in Gemma, a total of 54316 terms have been used to annotate their “topics”; 97% of which (52762/54316) were used to annotate human, mouse, and rat datasets - which we will further describe. Due to use of ontologies during curation, 98% (51479/52762) of dataset topics are represented by 9379 distinct ontology terms and the remainder are free-text terms (836 distinct free-text terms). The mean number of topics per dataset is 5.2.

In Gemma, the displayed dataset topics originate from three different levels of curation: dataset, experimental design, and sample-level curation. Dataset-level annotations are used to describe the general intent of the dataset and background information not captured at the other levels (e.g. “Alzheimer’s disease” for mouse models of Alzheimer’s, etc.); they are appended to the datasets directly, and represents 38% (20332/52762) of all dataset topics. Design-level annotations are used to describe the underlying experimental design of the study (i.e. different “factors” or sample groupings), while sample-level annotations are used to describe various sample characteristics. As design and sample-level annotations are not appended to the dataset directly, relevant annotations are algorithmically selected to reflect key information of the dataset, and are presented alongside dataset annotations as dataset topics. Design and sample-level annotations represent 56% (29393/52762) and 6% (3037/52762) of all dataset topics.

Separately, we also track the source of dataset annotations by adopting a framework derived from GO evidence codes; this reflects the two main methods in which annotations are appended to various Gemma entities: manual curation and automation (see Methods). The bulk of dataset topics are manually curated (code “IC” or “inferred by curator”), representing 95% of all dataset topics (49890/52762); while the remainder are automated (code “IEA” and “IIA”, or “inferred from electronic annotation” and “inferred from imported annotation” respectively). We next describe summaries of dataset topics using three different groupings (reflective of the core ontologies used): tissue/cell-type, disease, and chemical substances.

Of the 10420 human, mouse, and rat datasets, 80% (N = 8292) of them contain a dataset topic annotation related to tissues or cell types (i.e. usage of ontology terms from the UBERON or CL ontologies). Collectively, these topic annotations represent 29% (15202/52762) of dataset topics. After ontology inference, the number of “dataset-to-topic” associations increases to 265,087 in total. After manual inspection, we show the top 10 terms of varying resolutions that describe the variety of tissues and cell types represented in Gemma’s datasets (Figure 5). In Figure 5A, 34% (3517/10420) of datasets are associated with the “nervous system” [UBERON_0001016] - a reflection of our focused curation on nervous system related datasets in recent years. This is also reflected at the organ/tissue-level (Figure 5B) in which “brain” [UBERON_0000955] and “spinal cord” [UBERON_0002240] datasets collectively represent 25% (2651/10420) of Gemma datasets. In a similar vein, datasets related to the “hemolyphoid system” [UBERON_0002193] (Figure 5A), when grouped at the cell type resolution, consist of different “subtopics” (i.e. leukocyte, macrophage, T cell and B cell; Figure 5C).

**Figure 5:**
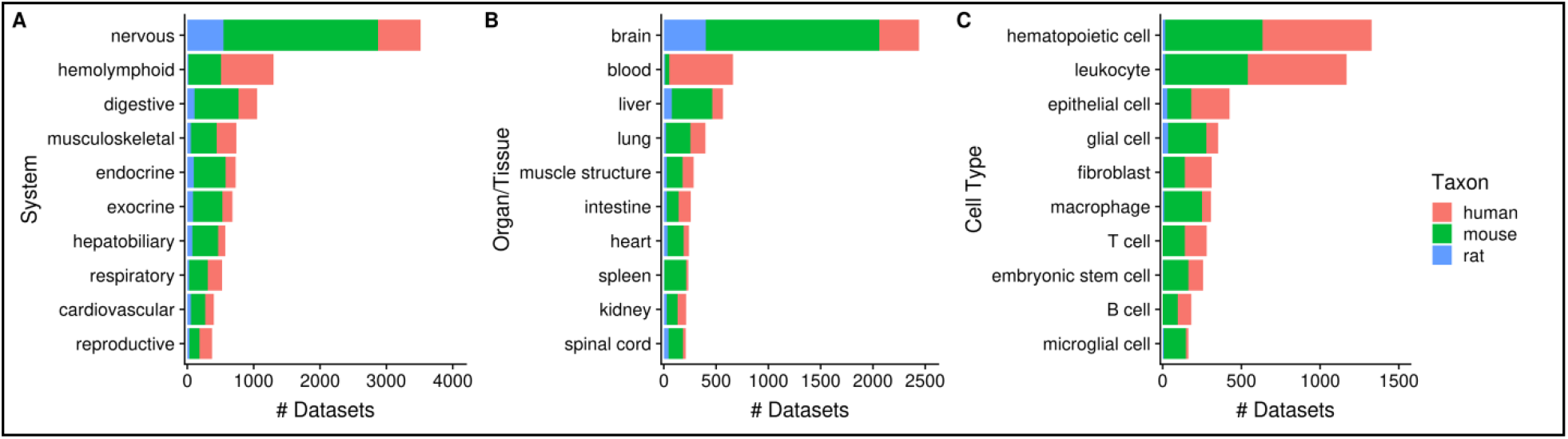
Numbers of datasets grouped by body system (A), organ/tissue (B), and cell type (C). The values are further grouped by taxon, and are represented using different colours. (N_dataset_ = 8292).

As mentioned earlier, by annotating datasets with ontology terms, we can utilize ontology inference to improve dataset retrieval during searches by expanding queries to child terms. In Figure 6, we show that the potential improvement in dataset retrieval is substantial for more general terms that have many children in the ontology. For example, the number of “brain”-associated datasets retrieved increases from 560 to 2442 with ontology inference, a 4.4-fold increase (Figure 6A); other terms with major improvements include “muscle structure” and “intestine”. We observe similar improvements at the cell type level, for terms such as “hematopoietic cell” (from 6 to 1328; 221-fold increase), “leukocyte”, “epithelial cell” etc. More specific terms naturally do not benefit as much from inference, e.g. “spleen” and “liver” (Figure 6A); and some do not increase at all, e.g. “embryonic stem cell” and “microglial cell” (Figure 6B). Again, this is reflective of the terms’ specificity: while “microglial cell” has two children terms, “brain” has 1974 of them.

**Figure 6:**
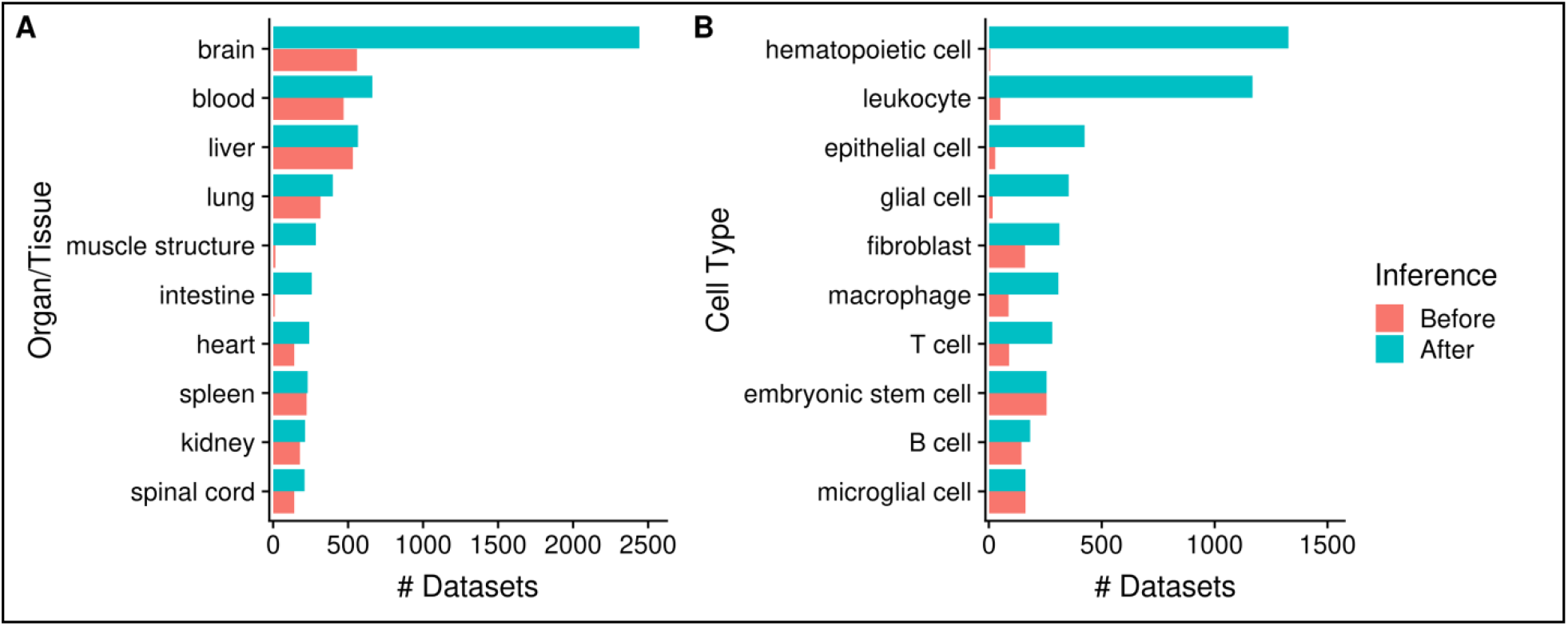
Numbers of datasets grouped by organ/tissue (A), and cell type (B). The colours indicate the number of associated datasets before (red) and after (blue) ontology inference. (N_dataset_ = 8292).

Next, 35% (3602/10420) of human, mouse, and rat datasets contain a dataset topic annotation related to diseases and disorders (i.e. usage of ontology terms from the DO ontology). These topic annotations represent 9% (4849/52762) of dataset topics, with the number of topic-dataset associations increasing to 36,797 after inference. In Figure 7A, we see that 17% (1733/10420) of datasets are associated with “cancer” [DOID_I62], followed by “nervous system disease” [DOID_863] (9%; 903/10420). Further inspection of ten “cancer” children terms showed that human datasets constitute an overwhelming majority of cancer-related datasets (Figure 7B), unlike the more “mouse-heavy” distribution observed in Figure 5 (i.e. tissue/cell-type topics). In the case of neuronal disorders, a more varied taxa-distribution is observed (Figure 7C). Disorders such as schizophrenia, bipolar disorder and major depressive disorder are overwhelmingly represented by human datasets, while diseases such as Alzheimer’s, amyotrophic lateral sclerosis and Huntington’s are more “mouse-heavy”; in the case of epilepsy datasets, “rat-heavy”. This is likely a reflection of the heavier dependence on animal models in research for certain disorders, and postmortem human brain tissue in others.

**Figure 7:**
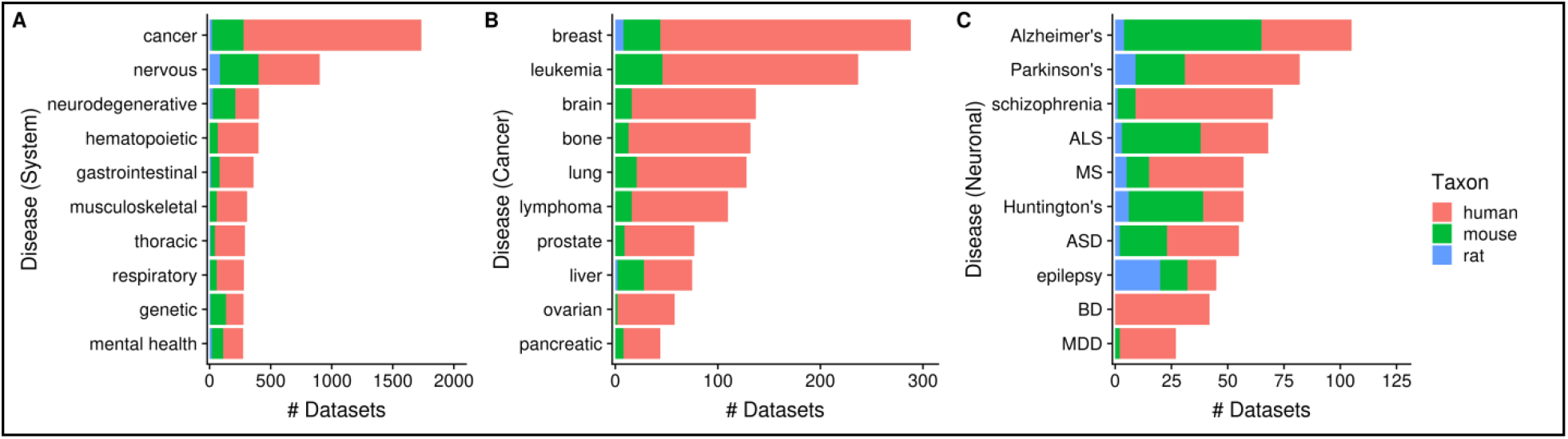
Numbers of datasets grouped by system-level classification of disorders (A), various cancers (B), and neuronal disorders (C). The values are further grouped by taxon, and are represented using different colours. (N_dataset_ = 3602). Abbreviations: amyotrophic lateral sclerosis (ALS), multiple sclerosis (MS), autism spectrum disorder (ASD), bipolar disorder (BD), and major depressive disorder (MDD).

Next, 23% (2362/10420) of datasets contain a topic annotation related to small molecules (i.e. terms from the ChEBI ontology). These annotations represent 7% (3837/52762) of dataset topics, with the number of topic-dataset associations increasing to 15,229 after inference. Figure 8A shows the ten most frequently used ChEBI terms in Gemma. The most frequently annotated ChEBI term, “lipopolysaccharide” [CHEBI_16412] (5%, 179/3837), is a compound found in the outer membrane of Gram-negative bacteria, and commonly used to study inflammatory responses and immune system function (e.g. in datasets GSE3253, GSE9509 and GSE15721; (52–54)). On the other hand, “ethanol” [CHEBI_I6236], the second most frequently annotated term (2%, 73/3837), is often used in modelling Fetal Alcohol Spectrum Disorder (e.g. GSE23115 and GSE18162; (55,56)) or study alcoholism in general (e.g. GSE13524 and GSE15774; (57,58)).

**Figure 8:**
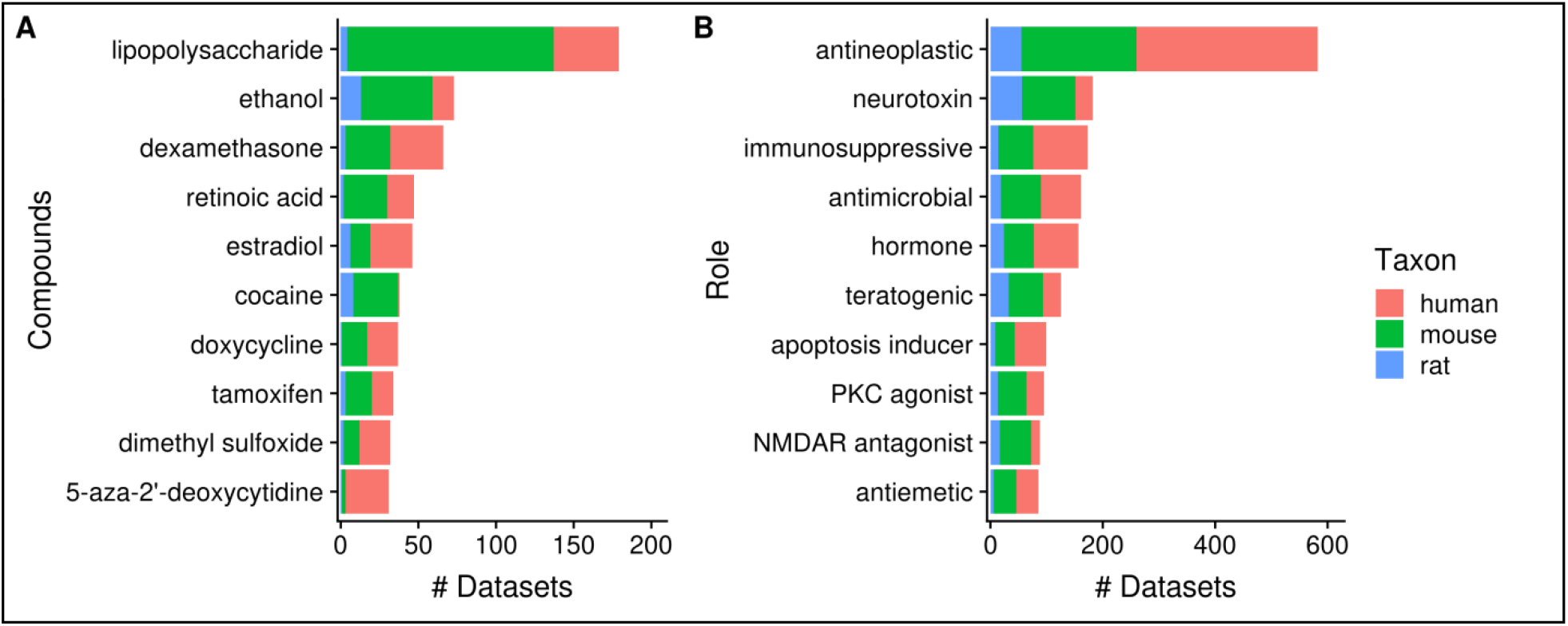
Numbers of datasets grouped by chemical compounds (A) and biological role/application of compounds (B). The values are further grouped by taxon, and are represented using different colours. (N_dataset_ = 2362).

As a further exploration, we summarized the annotation counts of compounds using children terms from two “role” branches of ChEBI: “biological role” [CHEBI_24432] and “application” [CHEBI_33232]; a selection of ten terms is shown in Figure 8B. The role term with the most number of associated datasets is “antineoplastic agent” [CHEBI_35610] (N = 582); the inferred associations deriving from 180 children terms used, including “dexamethasone” [CHEBI_41879] (N = 66) and “tamoxifen” [CHEBI_41774] (N = 34), both of which are listed in Figure 8A. This is followed by “neurotoxin” [CHEBI_50910] that is associated with 182 datasets; 26 contributing children terms, including “ethanol”. Another example is “immunosuppressive agent” [CHEBI_35705] (N = 173), deriving its associations from 25 children terms, such as “dexamethasone”. One interesting observation is “doxorubicin” [CHEBI_28748] (N = 29), a widely used chemotherapeutic was not included under the term “antineoplastic agent” (59), indicating incompleteness of relationship annotation in ChEBI, a common problem in ontologies in general (60).

### User Interface

In this section, we describe a selection of information pages that are publicly accessible through the Gemma web interface. One of the most important is the dataset page (Figure 9) and its associated tabs (Figure 10). We use GSE8030 (https://tinyurl.com/Gemma-GSE8030; (61)) and GSE2426 (https://tinyurl.com/Gemma-GSE2426; (62)) as examples. On this page, basic information such as the taxon, number of samples and the number of expression vectors (i.e. platform elements, or “Profiles”) is shown. There are five value-added components we want to focus upon:

**Figure 9:**
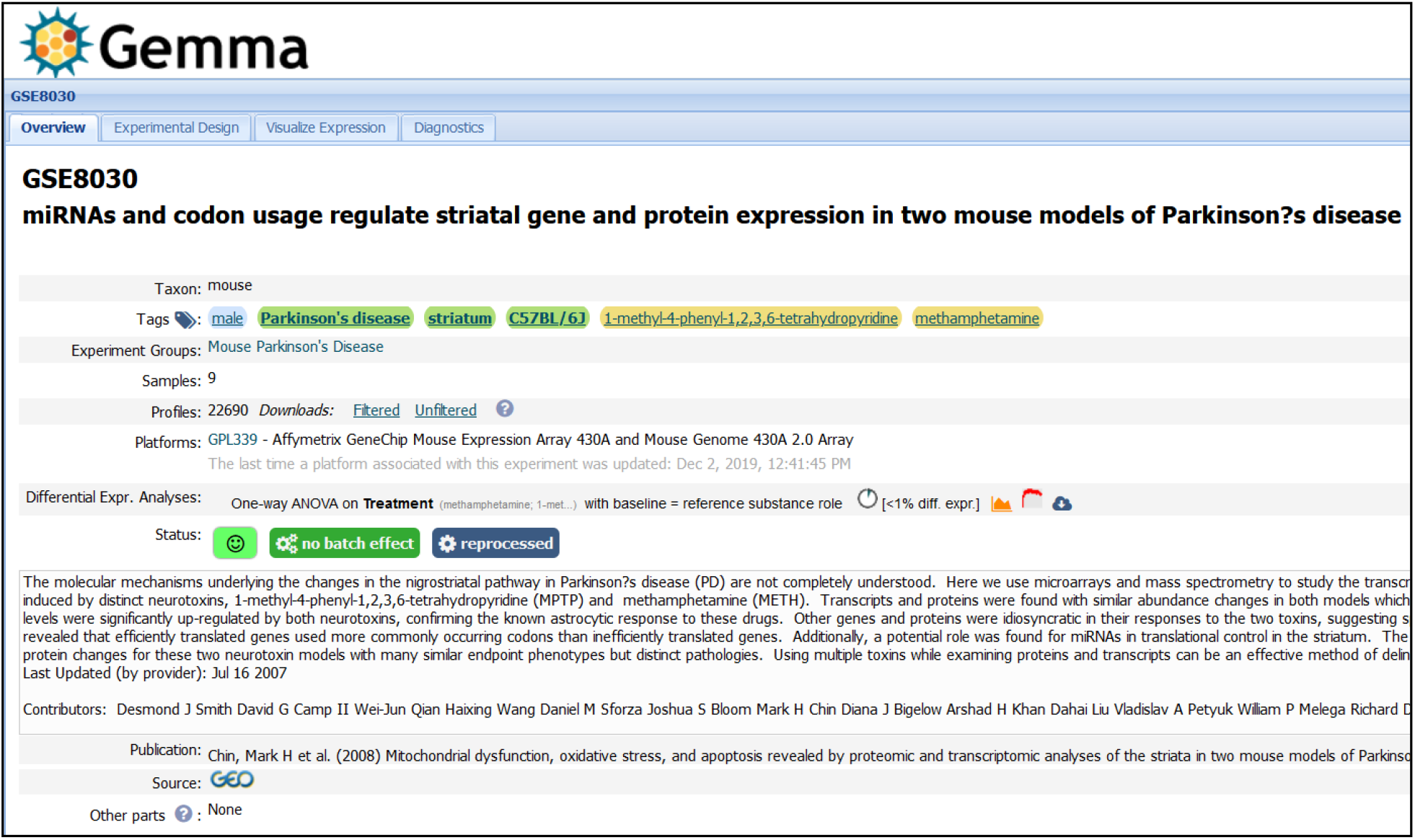
Snapshot of Gemma’s dataset page for GSE8030 (https://tinyurl.com/Gemma-GSE8030, (61)); see main text for description.

#### Dataset tags

We see there are six dataset tags appended to GSE8030 (e.g. male, Parkinson’s disease, etc.) These tags have three different background colours: green, yellow and blue. Green tags indicate the tag was added at the dataset level, yellow tags at the experimental design level (i.e. levels of a factor), and blue tags at the sample level. Both the design and sample level (i.e. yellow and blue) tags are propagated automatically to the dataset page. While the dataset and design level tags are added by manual curation, the sample level tags are generated automatically based on the parsed information obtain from GEO SOFT files.

#### Dataset status

A mix of colour-coded emoticons and visuals are used to display the state and quality of the dataset. Here, we see a green-smiley which represents a high GEEQ score. We also display batching information (in this case, GSE8030 has batch information and does not have batch effects), and data reanalysis state (GSE8030 has Affymetrix CEL files and was successfully reprocessed).

#### Differential expression analysis

Different visualizations are used to present the level of differential expression observed in each dataset (Figure 10C). The user can choose to view the distribution of p-values for each factor or a heat-map showing the top differentially expressed platform elements. A complete table containing the differential expression values (i.e. log_2_ fold change, t-statistics and p-value) for all the platform elements can also be downloaded for further inspection.

**Figure 10:**
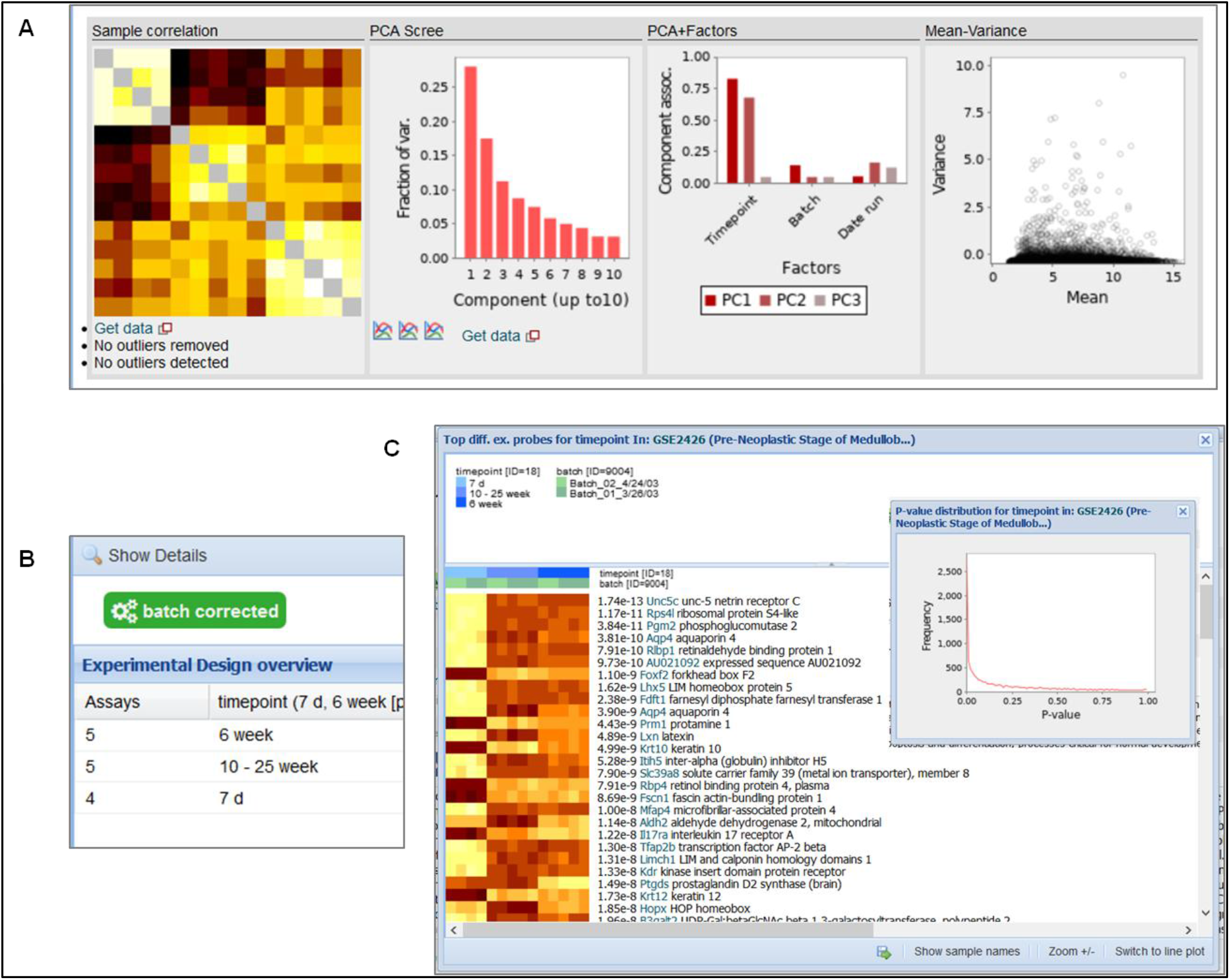
Snapshot of the contents in the “Diagnostics” tab (A); “Experimental Design” tab (B); and the “Differential Expression” buttons (C) for GSE2426 (https://tinyurl.com/Gemma-GSE2426, (62)). Within the diagnostics tab, a sample-sample gene expression correlation heat-map, PCA scree plot, PCA-Factor association plot and mean-variance scatterplot is displayed. For the differential expression details, a heat-map of the top differentially expressed platform elements and histogram of p-value distribution of the platform elements (inset) is provided.

#### Experimental design

Through the “Experimental Design” tab, the layout of the experimental design for the dataset is shown, with the number of samples in each combination of factor levels (Figure 10B). Additional columns are included for more factors; only the “batch” factor is not shown however.

#### Outlier removal

In the “Diagnostics” tab, we present different visualizations, including a heat-map of the sample-sample gene expression correlations, PCA scree-plots, bar plots showing the association of factors to the principal components, and a mean-variance scatter-plot (Figure 10A). Part of the manual quality control process involves removal of outlier samples, which can be observed as “greyed” out rows/columns in the sample correlation heat-map (the diagonal of the heat-map is also greyed out).

Another important page is the search page, where the end user could search for various entities, including genes, datasets (“Experiments”), and platforms that are included in Gemma. The example in Figure 11 shows a query for datasets that are annotated with the term “Parkinsons’s”.

**Figure 11:**
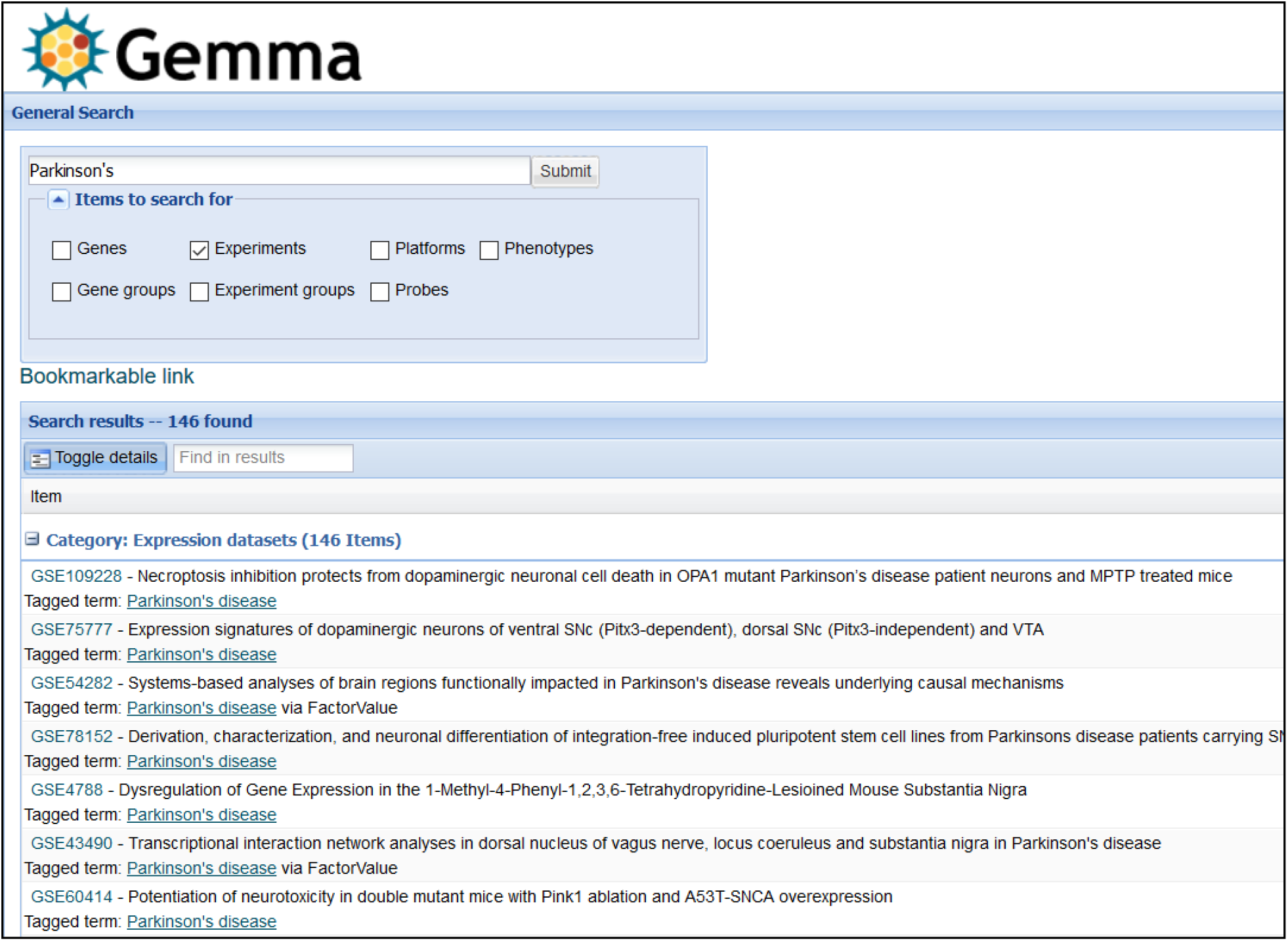
Snapshot of Gemma’s search page, in which datasets (“Experiments”) annotated with the term “Parkinson’s” is returned.

### Gemma REST API and R Package

For end users wishing to retrieve both data and metadata from Gemma programmatically, we implemented a REST-compliant API (Application Programming Interface) to assist in this task. Routine uses of this API include querying information on datasets, platforms and genes stored on the Gemma server. The output from these queries is structured as JSON files (Figure 12A). The complete documentation of available querying functions can be found here (https://tinyurl.com/Gemma-REST). For R users, we also provide an R package that utilizes the Gemma REST-API, returning the output as Robjects that is more convenient to use (https://github.com/PavlidisLab/gemmaAPI.R; Figure 12B); the documentation of this package is found here (https://pavlidislab.github.io/gemmaAPI.R/).

**Figure 12:**
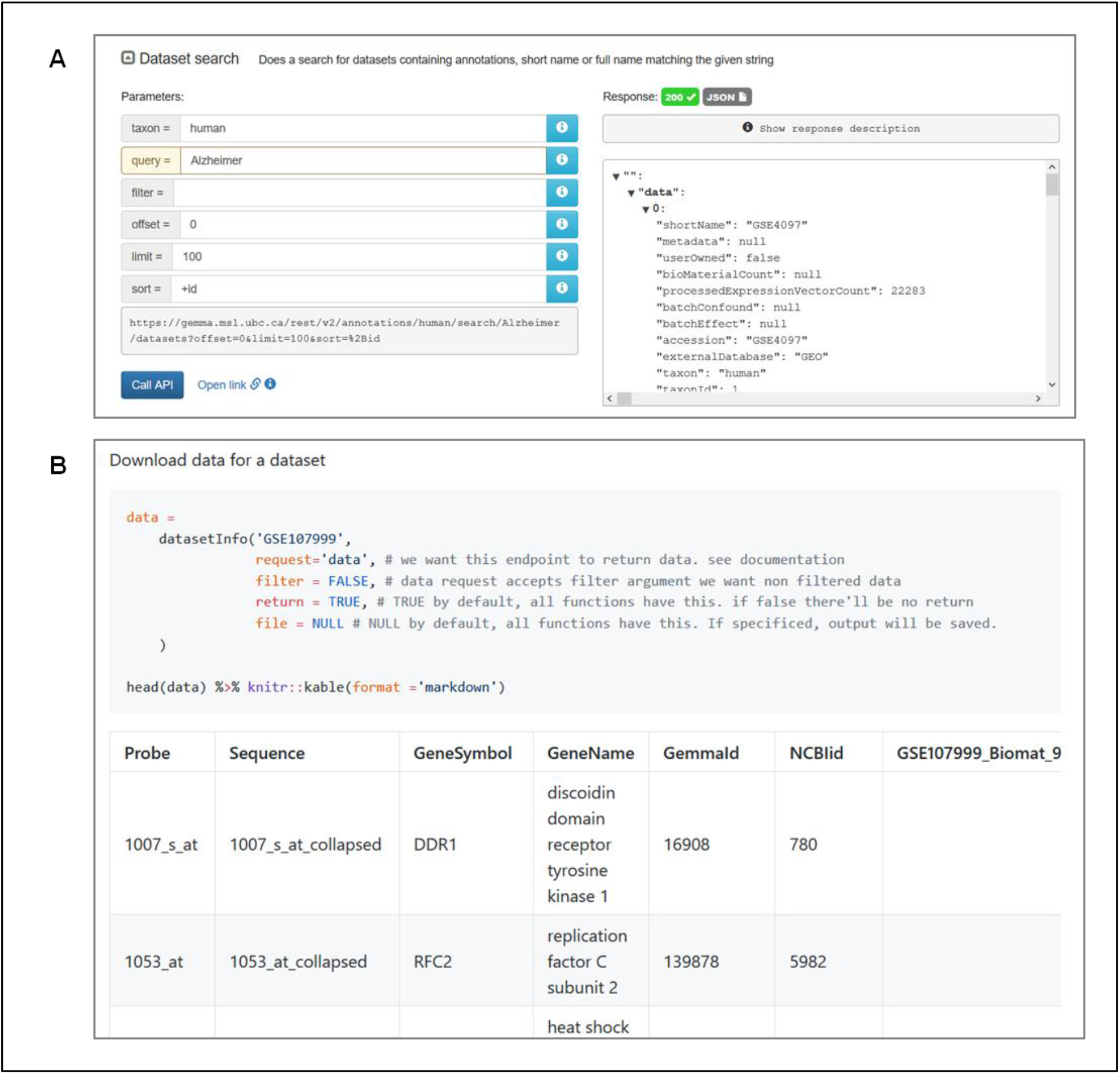
Snapshot of JSON output from searching for “Alzheimer” datasets using the interactive documentation website of the Gemma REST API (A); and tabulated output of GSE107999’s (63) metadata and expression data using the GemmaAPI.R package (B).

## Discussion

In this paper, we provide detailed and updated information on our efforts to facilitate use of publicly available gene expression profiling datasets. Using Gemma, researchers can circumvent the need to reprocess and reanalyze transcriptomic datasets, as all of that is performed by Gemma’s algorithms and our team of curators. Our curation efforts address some of the limitations of GEO’s data model. For microarray platforms, we remap probes to genes internally using an established protocol (20). Our efforts in curating experimental designs greatly facilitate use of the data. Problems relating to the data itself are addressed by our quality control checks, correction of batch artifacts whenever possible, removal of outliers, consistent processing of the expression data and Gemma’s internal mechanism of removing duplicate samples. Datasets considered unusable are blacklisted, and researchers can pre-emptively avoid them, saving valuable time and effort. Additional exploration and checks of the underlying gene expression data can also be performed using Gemma’s user interface.

To provide context of Gemma’s data holdings and processing pipelines, we compare Gemma to two other secondary gene expression databases: EMBL-EBI’s Gene Expression Atlas (64) and the Gene Perturbation Atlas (14). The Gene Expression Atlas was created to enable reuse of data residing in ArrayExpress (65), the European counterpart of the American NCBI GEO. As of June 2020, the Gene Expression Atlas (GXA) contains 3942 datasets covering 65 different taxa. Of those datasets, 64% (N = 2514) were imported from NCBI GEO; 32% (N = 802) of those GEO studies are also contained in Gemma. While GXA’s taxon coverage is fairly wide and would benefit researchers interested in taxa not included by Gemma, Gemma contains far more datasets; more so for human, mouse, and rat. Gemma’s more limited taxon coverage also enables us to focus our curation efforts on those three taxa. Another core difference is that GXA includes proteomics data, while Gemma solely processes transcriptomic data. Additionally, there are notable differences in the tooling and procedures for processing data. Batch correction of data is performed in Gemma using ComBat, while it is unclear if correction is implemented in GXA. For ontology support, while GXA relies exclusively on EFO (which does import terms from other ontologies), we rely on 12 different ontologies (including EFO), providing us greater coverage of concepts. The Gene Perturbation Atlas (GPA), on the other hand, was created to reuse expression data generated from human and mouse studies where genes were manipulated (e.g. knocked-down, overexpressed, etc.). For experiments involving the manipulation of protein-coding genes, this atlas contains data from 1749 GEO microarray datasets; 40% (N = 692) of which are also contained in Gemma. Due to GPA’s focus, users are constrained to genetic manipulation datasets, whereas Gemma provides the flexibility to also work with non-genetic manipulation datasets at the same time. Gemma also provides data from both microarray and RNA-seq studies, while GPA is currently solely microarray-based. Additionally, it is unclear whether the data in GPA was reprocessed uniformly, or if reprocessing from raw data was performed. Ontologies are not used in GPA, potentially limiting the potential for cross-database connectivity in the future.

Throughout this paper, we have emphasized Gemma’s utility in allowing access to data and analysis results for datasets in GEO. We have not detailed other analytical capabilities (co-expression and metaanalysis for instance) in Gemma, largely because these features are currently undergoing redesign and revision. As a data resource, Gemma’s main shortcoming is that we only have a fraction of the data contained in GEO. This is a gap that is unlikely to be closed, because the generation of data outstrips available resources to perform curation. As such, we have elected to focus our efforts in curating rodent and human datasets we deem relevant to neurodevelopment, neurological and neuropsychiatric conditions - a more manageable goal considering the annual submission rate of such datasets to GEO is approximately 500. However, we do continue to add data on other topics, and are able to respond to user requests for specific datasets.

There are several features that we plan to implement in Gemma. Single-cell RNA-sequencing is rapidly gaining wider adoption, and we anticipate a growing number of cell type-specific studies with biological replication, which we will be working to accommodate. Second, we will be enhancing the display and interpretation of differential expression on Gemma’s “Gene Information” pages. Recently, we showed that for certain genes, there is some level of predictability in their differential expression (66). This implies that genes that are often differentially expressed are less likely to be specific to a particular condition of interest, and we want to present this in our gene information panels. Another function of the “Gene Information” page is to provide further information on the conditions in which a gene is found differentially expressed, and can be used to infer the gene’s function. Our observation of pre-existing tools indicates that this information is often represented as a long list of “relevant conditions” (64). While manual inspection of these lists may provide some level of insight, it is time-consuming and unwieldy, especially for genes found to be differentially expressed in many conditions; and this issue will grow in severity alongside the continued increase of transcriptomic datasets. We are currently examining the feasibility of using ontology inference-based summarization techniques to improve usability of such lists.

## Acknowledgements

We are grateful for the efforts of past and present Gemma curators: Raymond Lim, Jenni Hantula, John Choi, Artemis Lai, Cathy Kwok, Celia Siu, Lucia Tseng, Lydia Xu, Mark Lee, Olivia Marais, Roland Au, Suzanne Lane, Tianna Koreman, Willie Kwok, Yiqi Chen, Tamryn Loo, Nathan Eveleigh, Fangwen Zhao, Nathan Holmes, Brenna Li, James Liu, Patrick Savage, Ellie Hogan, Dawson Born, Cindy-Lee Crichlow, Sophia Ly, John Phan, Brandon Huntington, Jimmy Liu, Aman Sharma, Calvin Chang, Henry Nguyen, Owen Tsai, Danja-Currie Olsen, Fiel Dimayacyac, and Morris Huang. We thank Jenni Hantula, Nathan Eveleigh and Raymond Lim for contributions to the development of the outlier detection algorithm. We also thank users from around the world who provided feedback and suggestions. We thank GEO and GEO data submitters for providing a rich source of data. We are also indebted to developers of the ontologies used.

## Author Contributions

P.P. designed and supervised the study; N.L. led curation efforts and designed research; N.L. and J.S. performed data analysis; P.P., S.T., M.B., M.J., and P.T. contributed to the Gemma software codebase; G.P.M. and M.B. assembled the RNA-sequencing processing pipeline; B.O.M. wrote the code for GemmaR; P.P. and N.L. wrote the manuscript.

